# Parsing the role of NSP1 in SARS-CoV-2 infection

**DOI:** 10.1101/2022.03.14.484208

**Authors:** Tal Fisher, Avi Gluck, Krishna Narayanan, Makoto Kuroda, Aharon Nachshon, Jason C. Hsu, Peter J. Halfmann, Yfat Yahalom-Ronen, Yaara Finkel, Michal Schwartz, Shay Weiss, Chien-Te K. Tseng, Tomer Israely, Nir Paran, Yoshihiro Kawaoka, Shinji Makino, Noam Stern-Ginossar

## Abstract

Severe acute respiratory syndrome coronavirus 2 (SARS-CoV-2) is the cause of the ongoing coronavirus disease 19 (COVID-19) pandemic. Despite its urgency, we still do not fully understand the molecular basis of SARS-CoV-2 pathogenesis and its ability to antagonize innate immune responses. SARS-CoV-2 leads to shutoff of cellular protein synthesis and over-expression of nsp1, a central shutoff factor in coronaviruses, inhibits cellular gene translation. However, the diverse molecular mechanisms nsp1 employs as well as its functional importance in infection are still unresolved. By overexpressing various nsp1 mutants and generating a SARS-CoV-2 mutant in which nsp1 does not bind ribosomes, we untangle the effects of nsp1. We uncover that nsp1, through inhibition of translation and induction of mRNA degradation, is the main driver of host shutoff during SARS-CoV-2 infection. Furthermore, we find the propagation of nsp1 mutant virus is inhibited specifically in cells with intact interferon (IFN) response as well as *in-vivo*, in infected hamsters, and this attenuation is associated with stronger induction of type I IFN response. This illustrates that nsp1 shutoff activity has an essential role mainly in counteracting the IFN response. Overall, our results reveal the multifaceted approach nsp1 uses to shut off cellular protein synthesis and uncover the central role it plays in SARS-CoV-2 pathogenesis, explicitly through blockage of the IFN response.

## Introduction

Severe acute respiratory syndrome coronavirus 2 (SARS-CoV-2) is the cause of the ongoing coronavirus disease 19 (COVID-19) pandemic (Zhu et al., 2020). COVID-19 presents a spectrum of clinical manifestations, ranging from asymptomatic infection to severe pneumonia. What determines the severity of COVID-19 is still not fully understood, but an imbalanced immune response seems to contribute substantially (King and Sprent, 2021).

Upon infection, IFN production is rapidly triggered by a series of host sensors that detect the presence of viral RNAs, leading to transcription and secretion of type I IFNs(Streicher and Jouvenet, 2019). Binding of IFN to its cognate receptor in autocrine and paracrine manners leads to the propagation of the signal and to the expression of hundreds of IFN stimulated genes (ISGs) that act to hamper viral replication at various stages of the viral life cycle (Schoggins, 2019). In the case of SARS-CoV-2, increasing evidence suggests that the lack of an effective type I IFN response plays a critical role in pathogenesis (Hadjadj et al., 2020; Hartenian et al., 2020; Meffre and Iwasaki, 2020; Sa Ribero et al., 2020).

Viruses have evolved diverse means to antagonize the IFN response. Specifically, SARS-CoV-2 encodes at least ten proteins whose over-expression counteracts the induction and antiviral action of IFNs(Sa Ribero et al., 2020). In addition to immune evasion, all viruses rely on the cellular ribosomes to translate their mRNAs into polypeptides. A common viral strategy to achieve this is to impair the translation of cellular mRNAs in a process termed “host shutoff”. Host shutoff is thought to contribute to viral takeover in two main ways: It helps to redirect translational resources and specifically ribosomes toward viral mRNAs and it blocks the ability of infected cells to mount an efficient antiviral response(Stern-Ginossar et al., 2019).

Viruses utilize diverse strategies to shutoff host protein synthesis, including interference in: mRNA transcription; processing and export out of the nucleus; and induction of mRNA degradation, as well as inhibition of the translation machinery (Stern-Ginossar et al., 2019). Coronaviruses (CoVs) are known to cause host shutoff, and several strategies have been proposed for how CoVs shutoff host protein synthesis (Abernathy and Glaunsinger, 2015; Nakagawa and Makino, 2021; Nakagawa et al., 2016). We recently demonstrated that SARS-CoV-2 infection leads to a reduction in translation capacity, degradation of host mRNAs, and repression of nuclear mRNA export (Finkel et al., 2021a). Although several viral proteins were suggested to play a role in SARS-CoV-2 induced shutoff (Banerjee et al., 2020; Lei et al., 2020; Miorin et al., 2020; Shemesh et al., 2021; Vazquez et al., 2021), nsp1 is the best characterized shutoff factor. Furthermore, studies with mutants lacking nsp1 in mouse hepatitis virus (MHV) and in SARS-CoV illustrate nsp1 is a major pathogenesis factor of beta CoVs (Jimenez-Guardeño et al., 2015; Shen et al., 2019; Züst et al., 2007)

Fundamental early work on SARS-CoV nsp1 (Huang et al., 2011; Kamitani et al., 2006; Lokugamage et al., 2012; Narayanan et al., 2008) together with recent studies on SARS-CoV-2 nsp1 show it interferes with host protein synthesis through several potentially distinct molecular mechanisms. Nsp1 binds directly to the 40S ribosomal subunit, and its C-terminal domain physically blocks the mRNA entry channel, leading to reduction in translation both in vitro and in cells (Lapointe et al., 2021; Schubert et al., 2020; Thoms et al., 2020; Tidu et al., 2020; Yuan et al., 2020). However, the extent by which the virus overcomes this nsp1-mediated translational block and how nsp1’s effects on translation come into play during infection remain largely unclear. In addition to blocking translation, SARS-CoV-2 nsp1 also promotes cellular mRNA degradation and viral transcripts are refractory to these effects through their 5’ leader sequence(Finkel et al., 2021a; Mendez et al., 2021). Finally, nsp1 was also shown to interact with the export factor NXF1 and to inhibit the nuclear export of cellular transcripts(Zhang et al., 2021). These findings portray complex interactions of nsp1 with the post-transcriptional life of mRNAs. Since there is extensive cross-talk between export, translation and degradation, and perturbation of one can lead to indirect effects on the others, it remains critical to evaluate the extent by which nsp1 directly perturbs each of these cellular functions and to measure the contribution of each function to the total shutoff effect. Furthermore, since various other viral proteins were suggested to interfere with cellular gene expression(Addetia et al., 2021; Banerjee et al., 2020; Hsu et al., 2021) and since host shutoff is thought to serve two independent purposes (to increase viral transcript translation and to inhibit the antiviral response), deciphering the role nsp1 plays in host shutoff in the context of infection and untangling its functional significance represent a critical knowledge gap.

Here we use overexpression of various SARS-CoV-2 nsp1 mutants to measure nsp1’s effects on cellular gene expression. We illustrate that nsp1 mediates its function by accelerating mRNA degradation, inhibiting translation and interfering with nuclear mRNA export, and we show that these mechanisms are likely distinct and complementary. We further generated a SARS-CoV-2 mutant in which nsp1’s ability to bind to the ribosome and to degrade cellular mRNAs is abolished. Using this mutant, we show that nsp1 is the main shutoff factor of SARS-CoV-2. Furthermore, we demonstrate that relative to cellular genes, nsp1 enhances the translation of viral genes. However, since nsp1 profoundly reduces the overall translation capacity in infected cells, in absolute terms (not relative to cellular genes), viral gene expression in cells infected with an nsp1 mutant virus is intact. Finally, we reveal the propagation of the mutant virus is inhibited both *in-vitro*, in cells with functional IFN response, and *in-vivo*, in infected hamsters and this attenuation is due to stronger IFN response. These results illustrate the multifaceted approach nsp1 uses to shut off cellular protein synthesis and the critical and specific role nsp1 plays in blocking the type I IFN response during SARS-CoV-2 infection.

## Results

### Nsp1 promotes both translation inhibition and decay of cellular mRNAs

We have previously demonstrated that in addition to SARS-CoV-2 nsp1’s ability to inhibit translation(Lapointe et al., 2021; Schubert et al., 2020; Shi et al., 2020; Thoms et al., 2020; Tidu et al., 2020), it also promotes RNA degradation (Finkel et al., 2021a; Mendez et al., 2021). In SARS-CoV nsp1, ribosome binding as well as residues R124/K125 are required for nsp1-induced host mRNA degradation activity (Kamitani et al., 2009; Lokugamage et al., 2012). Since nsp1 from SARS-CoV and SARS-CoV-2 are highly similar (84% identity in amino acids), we examined whether these residues are necessary for mRNA degradation also in SARS-CoV-2. We generated two nsp1 mutants. In the first mutant, nsp1-ΔRB (RB stands for ribosome binding), we removed amino acids 155-165, that are located in the C-terminal region of nsp1 and are critical for nsp1’s interactions with the ribosome (Schubert et al., 2020; Thoms et al., 2020; Tidu et al., 2020). In the second, nsp1-CD (CD stands for cleavage defective), we mutated R124 and K125 to alanine (R124A/K125A)(Figure 1a). We show that nsp1-WT, but not nsp1-ΔRB, associates with ribosomes (figure 1b), as previously reported (Mendez et al., 2021; Schubert et al., 2020; Thoms et al., 2020). In agreement with recent results (Mendez et al., 2021), nsp1-CD is still associated with ribosomes but to a lesser extent compared to nsp1-WT (Figure 1b). Notably, we noticed that nsp1 also likely represses its own expression, as the level of nsp1-WT protein was lower compared to the mutants that are likely functionally defective (Figure 1b and extended figure 1a). To test the inhibitory effects of these nsp1 mutants, we examined the expression of two GFP reporters, one in which the GFP is fused to human beta-globin 5’ UTR (host-5’UTR-GFP) and a second in which the GFP is fused to the viral 5’ Leader sequence (CoV2-leader-GFP). These reporters were co-transfected with vectors encoding nsp1-WT, nsp1-ΔRB, nsp1-CD or a control plasmid. Nsp1-WT and to a lesser extent nsp1-CD suppress the expression of host-5’UTR-GFP, whereas nsp1-ΔRB had no major effect on GFP expression (Figure 1c). The GFP reduction mediated by nsp1-WT was associated with ~6 fold reduction in GFP mRNA levels, whereas the levels of GFP mRNA were not affected by nsp1-CD or nsp1-ΔRB expression (Figure 1d). These results illustrate that as was reported for SARS-CoV nsp1(Kamitani et al., 2009; Lokugamage et al., 2012), binding to the ribosome is critical for promoting RNA degradation. Furthermore, since nsp1-CD maintains the ability to suppress cellular RNA translation, albeit to a lesser extent than that of nsp1-WT, but does not lead to mRNA degradation, these results indicate that the induction of RNA degradation and translation inhibition represent two distinct activities. In contrast to host-5’UTR-GFP and in agreement with previous findings (Finkel et al., 2021a; Mendez et al., 2021; Tidu et al., 2020), nsp1-WT did not inhibit, and even slightly increased, the level of CoV2-leader-GFP. Interestingly, in agreement with a recent publication (Mendez et al., 2021), nsp1-CD represses GFP expression even when it is fused to the CoV2-leader. Finally, as expected, nsp1-ΔRB had no major impact on CoV2-leader-GFP expression (Figure 1e). Analysis of RNA expression demonstrated that CoV2-leader-GFP was refractory to nsp1-WT induced degradation and similarly to host-5’UTR-GFP both nsp1-CD and nsp1-ΔRB did not drastically affect CoV2-leader-GFP RNA expression (Figure 1f).

**Figure 1:**
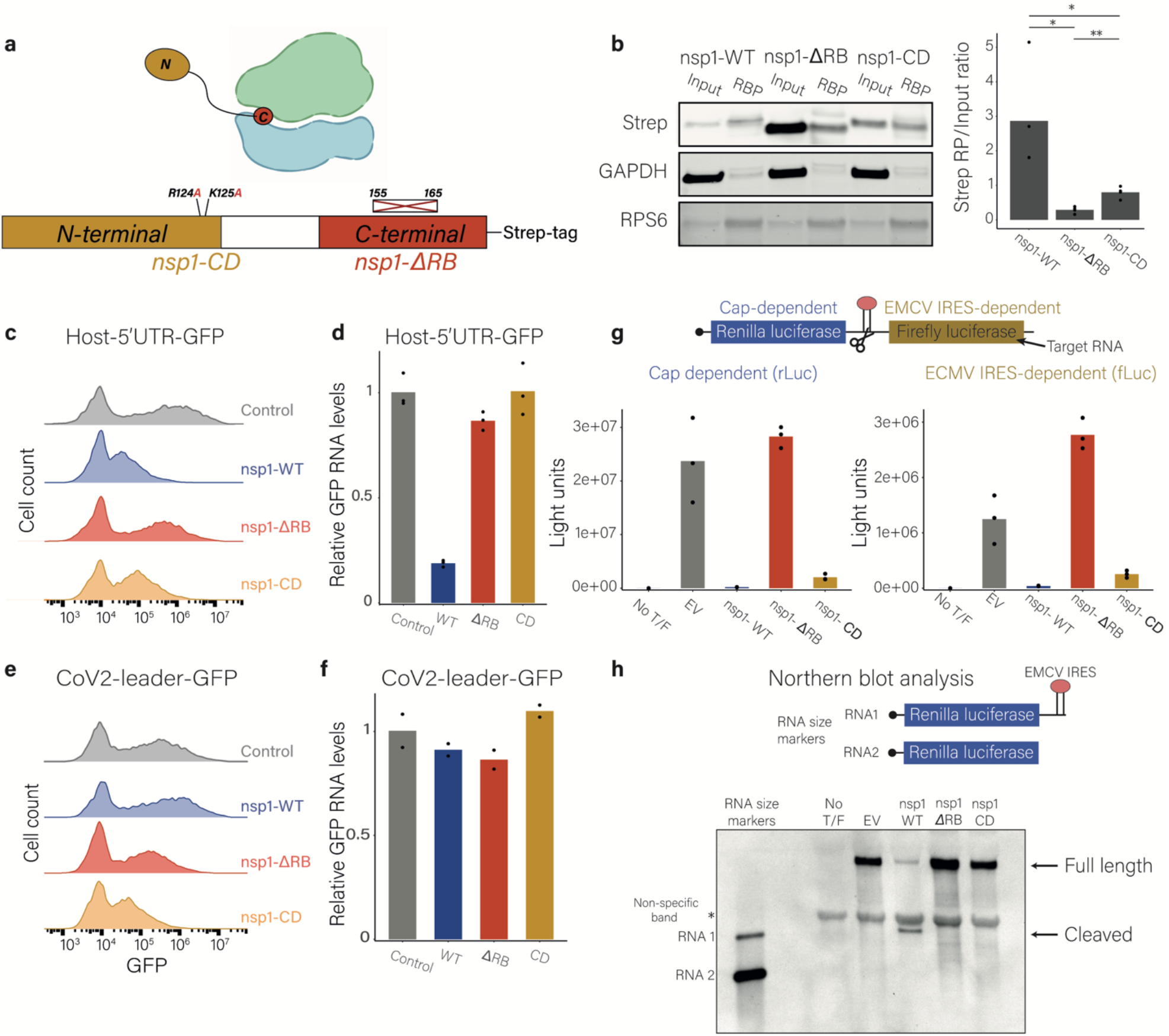
Nsp1 promotes both translation inhibition and decay of cellular mRNAs. **a**, Schematic illustration of the nsp1-ΔRB and nsp1-CD mutation. **b**, 293T cells were transfected with plasmids expressing nsp1-WT, nsp1-ΔRB, or nsp1-CD, and ribosomes were pelleted. Nsp1 protein levels in the input and in the ribosome pellet (RBP) were measured by Western blot with Strep-Tactin. RPS6 and GAPDH were used as ribosomal protein and loading controls, respectively. Right panel presents the ratio of the strep signal in the RBP to input, which was assessed using the intensity of the bends. Points show technical replicates, * = *p*<0.05 and ** = *p*<0.01 for a two tailed t-test. **c-f**, 293T cells co-transfected with plasmids expressing nsp1-WT, nsp1-ΔRB, or nsp1-CD or a control plasmid together with (**c** and **d**) a reporter plasmid expressing GFP fused to human beta-globin 5’UTR (host-5’UTR-GFP), or (**e** and **f**) the SARS-CoV-2 5’leader sequence (CoV2-leader-GFP). (**c** and **e**) GFP expression was measured by flow cytometry. (**d** and **f**) Relative GFP mRNA levels were measured by quantitative real-time PCR. 18S ribosomal RNA was used as a normalizing gene. Points show measurement of technical replicates. (**g** and **h**) Vero E6 cells were co-transfected with the reporter plasmid containing a cap-dependent *Renilla* luciferase followed by an EMCV IRES and *Firefly* luciferase (pRL-EMCV-FL) along with pCAGGS empty vector (EV) as a control plasmid or a plasmid expressing nsp1-WT, nsp1-ΔRB or nsp1-CD. No T/F represents non-transfected cells. (g) Luciferase expression levels. Y-axis represents light units. (h) RNAs were extracted and analyzed by Northern blot using a digoxigenin-labeled rLuc riboprobe. RNA size marker is a mixture of in vitro-transcribed RNA transcripts, RNA 1 and RNA 2, as shown. RNA 1 contains the region from the 5’-end to the 3’-end of the intercistronic region of pRL-EMCV-FL, and RNA 2 contains the region from the 5’-end to the end of the rluc ORF in pRL-EMCV-FL. A non-specific bend is indicated.

We next examined how nsp1 affects translation that is mediated by an internal ribosome entry site (IRES). To this end, we used a bicistronic mRNA, in which expression of the upstream *Renilla* luciferase is mediated by cap-dependent translation and expression of downstream *firefly* luciferase is driven by EMCV IRES (Figure 1g). Nsp1-WT and to a lesser extent nsp1-CD but not nsp1-ΔRB suppressed the production of both *Renilla* and *Firefly* luciferases, indicating nsp1 inhibits both capped as well as IRES driven translation (Figure 1g). Furthermore, nsp1-WT but not nsp1-ΔRB or nsp1-CD induced endonucleolytic cleavage of the mRNA (Figure 1h). Overall, these results show that SARS-CoV-2 nsp1 blocks translation and induces endonucleolytic cleavage of cellular RNAs that leads likely to their degradation. Nsp1 binding to the ribosome is essential for both these activities but since not all translation block lead to RNA degradation these activities likely represent distinct functions.

### Nsp1 inhibits mRNA export independent of its ribosome binding activity

On top of the effects of nsp1 on translation and RNA degradation, nsp1 was demonstrated to block nuclear mRNA export through interaction with NXF1(Zhang et al., 2021), representing an additional mechanism through which nsp1 inhibits cellular gene expression. Notably, the interactions of nsp1 with NXF1 were suggested to be mediated by nsp1 N-terminal region (amino acids 1-129, figure 1a), and therefore are likely independent of nsp1 binding to the ribosome (Zhang et al., 2021). Indeed, using both host-5’UTR-GFP and CoV2-leader-GFP, nsp1-ΔRB still had a small effect on GFP expression compared to the control protein (1.5-fold and 1.3-fold relative reduction in MFl [median fluorescence intensity], respectively). To examine if nsp1 affects mRNA export and if this activity is maintained in the absence of ribosome binding, we transfected 293T cells with plasmid expressing nsp1-WT, nsp1-ΔRB or GFP (as a control) and assessed subcellular localization of polyadenylated transcripts by cytoplasmic/nuclear (cyto/nuc) fractionation followed by RNA-seq. Fractionation efficiencies were confirmed by comparing expression values of known nuclear and cytoplasmic RNAs as well as snoRNAs nuclear enrichment (extended figure 1b and c). In addition we obtained a strong correlation between our cyto/nuc measurements and measurements conducted previously in a different cell type (Finkel et al., 2021a) (extended figure 1d). In agreement with nsp1 inhibiting mRNA export, nsp1-WT expression led to relative nuclear enrichment of cellular transcripts (Figure 2a). Furthermore, this enrichment did not depend on nsp1 induction of mRNA degradation, as nsp1-ΔRB expression also led to relative nuclear enrichment, and the effects were even stronger than those of nsp1-WT. The stronger effect of nsp1-ΔRB was unexpected but could be attributed to the stronger expression of nsp1-ΔRB due to self-inhibitory effects of nsp1-WT (extended figure 1a). Indeed, analysis of nsp1 expression from the RNA-seq data showed that nsp1-ΔRB expression was 8-fold higher compared to nsp1-WT (extended figure 1e). Furthermore, it is possible that in the absence of binding to the ribosome more nsp1 was free to interact with NXF1. Altogether these measurements show that nsp1 is able to target three distinct steps in the host cell mRNA biogenesis pathway and that its ability to interfere in mRNA export does not depend on its binding to the ribosome.

**Figure 2:**
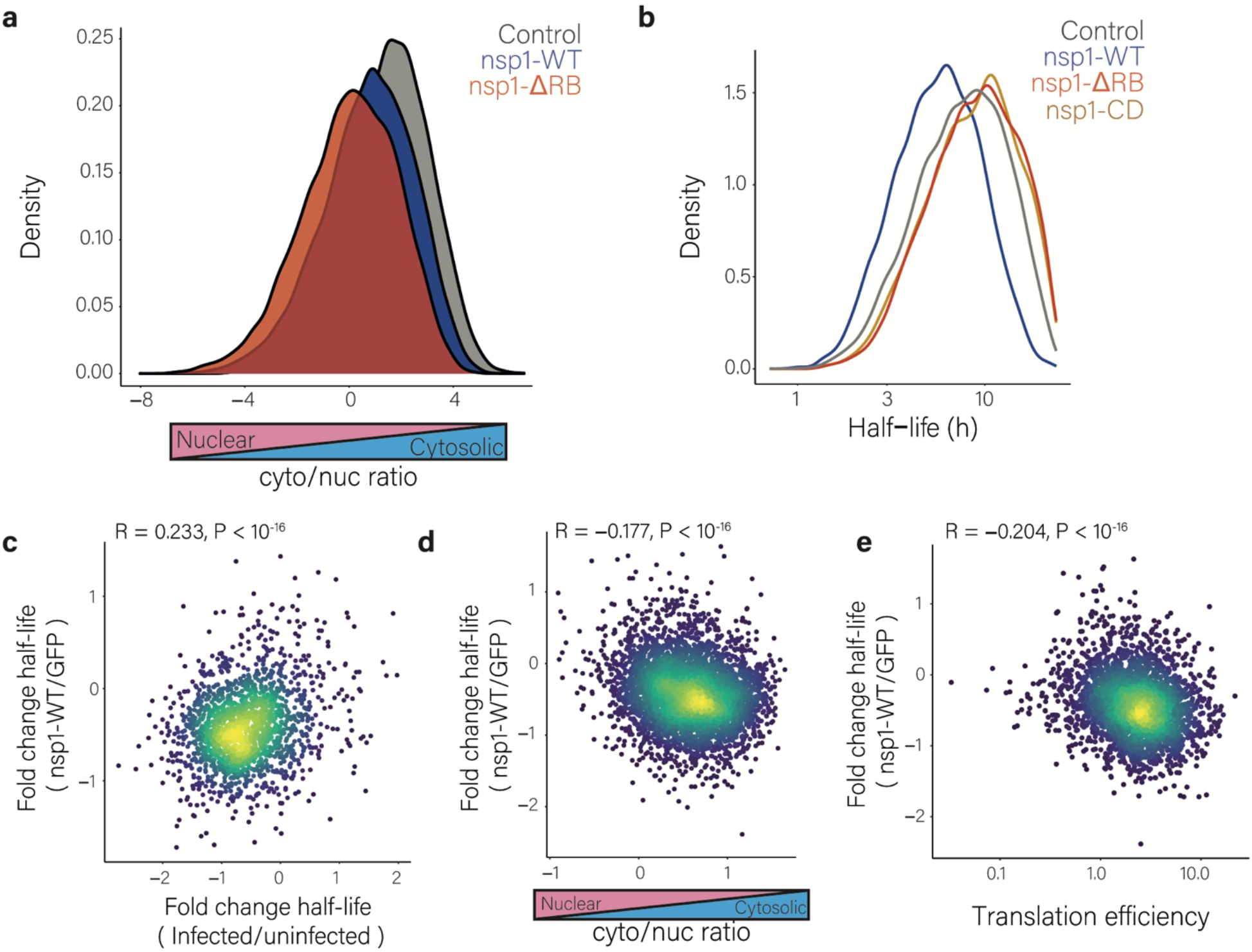
Nsp1 inhibits nuclear mRNA export and accelerates the decay of cytosolic mRNAs. **a**, Distribution of the cytosolic to nuclear ratio of cellular transcripts in 293T cells transfected with nsp1-WT, nsp1-ΔRB, or control expression plasmids that were subjected to subcellular fractionation followed by RNA-seq. **b**, Distribution of cellular transcript half-lives in 293T cells transfected with nsp1-WT, nsp1-ΔRB, nsp1-CD, or control expression plasmids. Half-lives were determined by SLAM-seq measurements followed by analysis using GRAND-SLAM (Jürges et al., 2018). **c**, Scatter plot of the fold change of transcript half-lives between SARS-CoV-2-infected cells and uninfected Calu3 cells (Finkel et al., 2021a), relative to the fold change of 293T cells transfected with nsp1-WT compared to a control plasmid. **d**, Scatter plot of the fold change of transcript half-lives between 293T cells transfected with nsp1-WT compared to control plasmid, relative to the cytosolic to nuclear ratio of the transcripts. **e**, Scatter plot of the fold change of transcript half-lives between 293T cells transfected with a plasmid expressing nsp1-WT or with a control plasmid, relative to their translation efficiency (TE) values from uninfected cells. Pearson’s *R* and two-sided *P* values are presented.

### Nsp1 induces degradation of cytosolic transcripts

We next wanted to characterize the cellular transcripts that are sensitive to nsp1-induced degradation in an unbiased manner. Since we observed significant differences in the expression of nsp1-WT, nsp1-ΔRB and nsp1-CD that are due to nsp1 self-inhibition, we fused the CoV2-5’ leader sequence upstream of the nsp1 constructs, to confer protection from self inhibition. In agreement with our GFP reporter results, nsp1-WT with the CoV2-leader was refractory to its self-inhibitory effects and its expression was even enhanced compared to the mutated constructs with the CoV-2 5’leader (extended Figure 2a), further illustrating that nsp1-WT promotes the translation of transcripts that contain the CoV2 5’leader.

To directly measure the induction of mRNA decay by nsp1 we employed SLAM-seq(Herzog et al., 2017). This approach allows measurement of endogenous mRNA half-lives based on 4-thiouridine (4sU) incorporation into newly transcribed RNA. After RNA extraction, 4sU is converted to a cytosine analogue using iodoacetamide, and these U to C conversions are identified and quantified by RNA sequencing, providing means to quantify old and new mRNAs and to calculate mRNA decay (Finkel et al., 2021a; Herzog et al., 2017; Jürges et al., 2018). We applied SLAM-seq to 293T cells transfected with plasmids expressing nsp1-WT, nsp1-CD, nsp1-ΔRB or GFP as control (Extended figure 2b). We obtained all characteristics of high-quality SLAM-seq libraries; >8000 quantified genes, U- to C-mutation rates starting at ~1.5% and rising to ~5.5% and an increase with time in the portion of labeled RNA, which was stronger in cells expressing nsp1-WT, indicating a faster turnover of RNA in these cells (extended figure 2c-d). There was strong correlation between half-lives estimated from our SLAM-seq measurements in the control cells (transfected with GFP) and previous half-lives measurements in a different cell type(Finkel et al., 2021a) (extended figure 2e), indicating the robustness of our measurements. Reassuringly, analysis of nsp1 expression from the RNA-seq reads demonstrated an equivalent expression of all constructs (extended figure 2f), illustrating that addition of the CoV2-5’ leader to nsp1 mRNA indeed prevented the nsp1 mRNA degradation. Importantly, we observed a substantial reduction in cellular mRNA half-lives in cells expressing nsp1-WT but not in cells expressing nsp1-ΔRB or nsp1-CD (Figure 2b), indicating induction of significant mRNA degradation only by nsp1-WT. Furthermore, although the measurements were conducted in different cell types, we observed significant correlation between the profile of mRNA degradation that is induced by SARS-CoV-2 infection(Finkel et al., 2021a) and the degradation induced by nsp1-WT but not by nsp1-ΔRB or nsp1-CD (figure 2c, extended figures 3a and 3b). This indicates that nsp1 plays a major role in inducing RNA degradation in infected cells and that this degradation is dependent on nsp1 ribosome binding. In accordance with our measurements in infected cells(Finkel et al., 2021a), expression of nsp1-WT, but not of nsp1-ΔRB or nsp1-CD, led to a more substantial reduction in the half-lives of cytoplasmic transcripts compared to transcripts that are mostly nuclear (Figure 2d and extended figures 3c and 3d). We also observed significant correlation between the translation efficiency of cellular genes and their half-life reduction following nsp1-WT expression (Figure 2e). We analyzed transcripts features that are associated with sensitivity to nsp1-induced degradation and the two features that best explain the sensitivity of mRNAs are cytosolic localization and higher translation efficiency (extended figure 3e). Overall, these findings demonstrate that induction of mRNA degradation is a distinct function of nsp1 which is independent of export inhibition, and is mainly directed at cytosolic mRNAs which are engaged with the ribosome.

**Figure 3:**
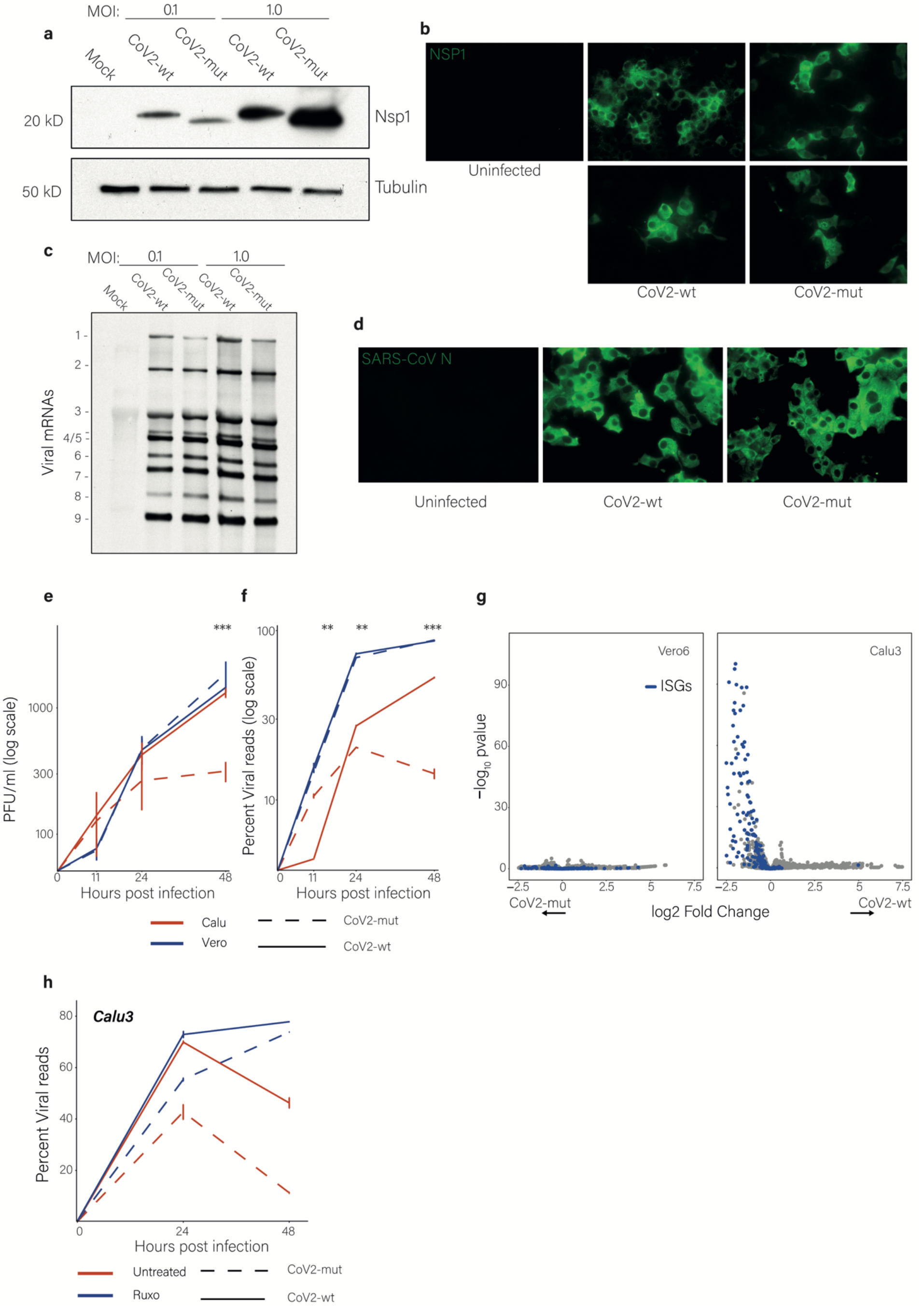
SARS-CoV-2 carrying nsp1-ΔRB is attenuated specifically in IFN competent cells. **a**, Western blot analysis of the Nsp1 protein in Vero cells infected with the CoV2-wt or with the CoV2-mut at an MOI of 0.1 or 1. **b**, Microscopy images of Vero cells infected with CoV2-wt or with CoV2-mut at 16 hpi with an MOI of 2, and stained with antibodies for Nsp1. **c**, Northern blot analysis using a probe, which binds to the common 3’ UTR of viral mRNAs and detects all viral mRNAs, in Vero cells infected with the CoV2-wt or with the CoV2-mut at MOI of 0.1 or 1 at 18hpi. **d**, Microscopy images of Vero cells infected with CoV2-wt or with CoV2-mut, and stained with antibodies for the nucleocapsid protein. (**e** and **f**) viral titers (**e**) or percentage of viral reads (**f**) in Calu3 or Vero cells infected with CoV2-wt or with CoV2-mut at 0, 11, 24, and 48 hpi with an MOI of 0.01. Two tailed T-test performed where ** = *p*<0.01 and *** = *p*<0.001. **g**, Volcano plots showing changes in cellular transcript levels in CoV2-wt versus CoV2-mut infected Calu3 (right) or Vero (left) cells at 24 hpi. The fold change and statistical significance are presented in the x axis and y axis, respectively. Enrichment of ISGs was calculated using a hypergeometric test. ISGs are marked in blue. **h**, percentage of viral reads in untreated or Ruxolitinib treated Calu3 cells infected with CoV2-wt or with CoV2-mut at 0, 24, and 48 hpi with an MOI of 0.01. The effect of Ruxolitinib treatment on CoV2-mut was significantly stronger compared to the effect on CoV2-wt, * = *p*<0.05 and *** = *p*<0.001 using linear regression.

### SARS-CoV-2 with a mutant nsp1 is attenuated in IFN competent cells

To evaluate the molecular and biological functions of nsp1 in infected cells, we generated a SARS-CoV-2 mutant, in which amino acids 155-165 of nsp1 were deleted by using a reverse genetics system (Xie et al., 2020) (CoV2-mut). This mutant is similar to the nsp1-ΔRB in the expression studies described above. Deep sequencing confirmed the introduced deletion (extended figure 4a), and there were no additional nucleotide alterations within the nsp1 gene or along the SARS-CoV-2 genome. Our initial characterization of the CoV2-mut was conducted in Vero cells (kidney epithelial cells from African green monkey) which are type I IFN-deficient(Emeny and Morgan, 1979). Cells infected with wild-type SARS-CoV-2 (CoV2-wt) or CoV2-mut produce similar levels of nsp1 protein (Figure 3a), confirming that the short deletion in nsp1 did not alter its expression. The majority of the nsp1-wt and nsp1-ΔRB accumulated in the cytoplasm in infected cells but nsp1-ΔRB also showed perinuclear localization (Figure 3b), potentially reflecting stronger engagement with nuclear export machinery in the absence of ribosome binding. Infection with CoV2-mut or CoV2-wt exhibited comparable levels of viral mRNAs at 18hpi (Figure 3c) as well as of the viral N protein (Figure 3d). Overall, these experiments illustrate that infection of Vero cells with CoV2-mut resulted in viral gene expression which is comparable to that of CoV2-wt.

**Figure 4:**
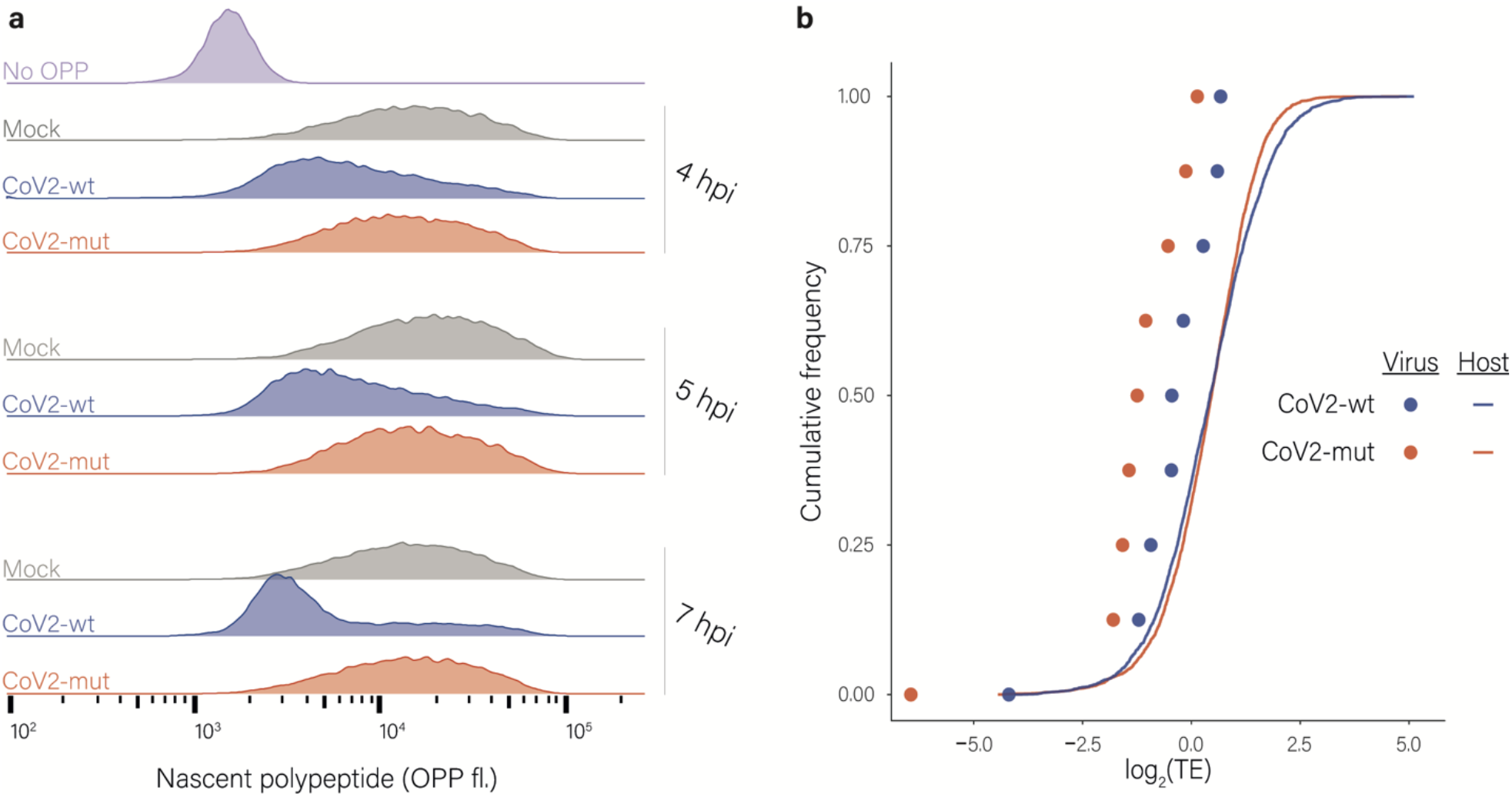
Effects of nsp1 on translation and accumulation of viral and host mRNAs. **a**, Protein synthesis measurement by flow cytometry of Calu3 cells infected with CoV2-wt or with CoV2-mut (MOI = 3) for 4, 5 and 7 hpi or an uninfected control following O-Propargyl Puromycin (OPP) incorporation and fluorescent labeling using Click chemistry **b**, Cumulative frequency of human (line) and viral (dots) genes according to their relative translation efficiency (TE) in cells infected with CoV2-wt (blue) or with CoV2-mut (red) at 4 hpi. TE was calculated from ribosome profiling and mRNA sequencing, and is defined as the ratio of ribosome footprints to mRNA for a given gene. Each dot represents one of nine major viral mRNA species.

We next analyzed the propagation of CoV2-mut and CoV2-wt in both Vero cells and Calu3 cells (lung adenocarcinoma cells), which have intact IFN-response. Both cells efficiently support SARS-CoV-2 replication (Finkel et al., 2021b). In Vero cells CoV2-mut titers were overall similar to CoV2-wt and at 48 hpi were even slightly higher than CoV2-wt (Figure 3e). In contrast, in Calu3 cells viral titers of CoV2-mut were lower compared to the CoV2-wt (at 48 hpi *p* = 0.00018 by t-test, Figure 3e) and this attenuation in CoV2-mut growth was also observed along a longer time course at a lower MOI (extended figure 4b). Consistent with the growth kinetics, analysis of viral gene expression using RNA-seq shows comparable levels of viral RNAs from the CoV2-wt and CoV2-mut along infection in Vero cells (Figure 3f). In contrast, in Calu3 cells, despite similar initial levels of viral RNAs, at 24 and 48 hpi viral gene expression was considerably lower for CoV2-mut compared to CoV2-wt (at 24 hpi *p* = 0.0016, and at 48 hpi *p* = 0.0004 by t-test, Figure 3f). The similar propagation of CoV2-mut and CoV2-wt in Vero cells versus the attenuation of CoV2-mut in Calu3 cells points that inhibition of viral propagation due to nsp1 mutation is due to host response to SARS-CoV-2 infection and not to direct effects of nsp1 on viral gene expression. Since the IFN response is critical to SARS-CoV-2 propagation (Felgenhauer et al., 2020; Kumar et al., 2021; Lei et al., 2020; Mantlo et al., 2020; Xia et al., 2020) and it is compromised in Vero cells (Emeny and Morgan, 1979) extended figure 4c), we speculated that these differences are due to nsp1’s interference, or lack thereof, with the IFN response. Indeed, analysis of changes in cellular gene expression showed a stronger induction of ISGs in Calu3 cells infected with CoV2-mut compared to those infected with CoV2-wt (*p* value = 1.55×10^−114^, Figure 3g). In contrast, no differences in the response to the two viruses were observed in infected Vero cells (Figure 3g). To confirm that the observed inhibition in viral growth is indeed due to a more potent IFN response, we tested if the inhibition of IFN signaling by Ruxolitinib (a selective JAK-STAT inhibitor (Jackson et al., 2016)), can rescue CoV2-mut propagation in Calu3 cells. Indeed, CoV2-mut propagation in Calu3 cells was rescued by Ruxolitinib treatment, whereas propagation in CoV2-wt was impacted to a reduced extent (Figure. 3h). We further confirmed that Ruxolitinib treatment abolished the differences in ISG expression between CoV2-mut and CoV2-wt infection (Extended Data Fig. 4d). Overall, these results illustrate that the altered activity of nsp1 impairs the propagation of SARS-CoV-2 due to an enhanced IFN response.

### Nsp1 ribosome binding supports viral RNA translation in infected cells

To gain insights into the molecular mechanisms that drive the functional differences between CoV2-wt and CoV2-mut, we assessed the changes that occur in viral and host translation and RNA expression during infection with these two viruses. We infected Calu3 cells with CoV2-wt or CoV2-mut at an MOI of 3, resulting in infection of the vast majority of the cells and thus a mostly synchronous cell population. In line with the early time points of the growth kinetics in Calu3 (figure 3e and 3f), at 8 hpi there were no substantial differences in the infection efficiency of the two viruses (extended data 5a), allowing for direct molecular dissection of the effects of CoV2-wt and CoV2-mut on cellular and viral gene expression.

We first examined how CoV2-wt and CoV2-mt affect global translation levels. To quantify absolute translation levels, we measured nascent protein synthesis levels using a tagged analogue of puromycin, O-Propargyl Puromycin (OPP), which is incorporated into elongating polypeptide chains((Finkel et al., 2021a) and extended data 5b). We infected Calu3 cells with CoV2-wt or CoV2-mut or mock infected, and measured nascent protein synthesis levels in mock infected cells and at 4, 5 and 7 hpi. CoV2-wt infection led to significant reduction in global translation levels already at 4 hpi which was augmented with time, and at 7 hpi translation activity was reduced by 4-fold (Figure 4a). In contrast, CoV2-mut had no major effect on the overall translation levels in infected cells (Figure 4a). These results illustrate that nsp1 is the main driver of translation shutoff during SARS-CoV-2 infection, and that this effect is mediated by its binding to the ribosome.

The OPP-labeling experiments provide an absolute measure of translation in infected cells, yet they do not indicate how much of the translation capacity is dedicated to the translation of viral genes. In order to gain a detailed view on the relative levels of viral and host translation, and to decipher the importance of nsp1 for viral RNA translation during infection, we infected Calu3 cells with with CoV2-wt or CoV2-mut and harvested infected cells at 4 and 7 hpi as well as uninfected cells for RNA-seq and ribosome profiling. Metagene analysis, in which gene profiles are aligned and then averaged, revealed the expected profiles of footprints and mRNAs (extended data 6a and 6b). Using this data, we quantitatively assessed the expression pattern of cellular and viral transcripts. In order to evaluate the ability of SARS-CoV-2 to co-opt the host ribosomes we calculated the translation efficiency (TE, ratio of footprints to mRNAs for a given gene) of viral and cellular mRNAs in infected cells. We then compared the TE of human genes to that of viral genes at each time point with each virus. The analysis shows that when cells are infected with CoV2-wt viral gene translation efficiencies fall within the low range of cellular gene translation (Figure 4b and extended figure 6c). This indicates that in cells infected with CoV2-wt, viral transcripts are translated with efficiencies that are slightly lower than those of host transcripts. This could be attributed to the sequestration of vast amounts of viral RNA in double-membrane replication compartments, rendering them inaccessible to the ribosomes (Wolff et al., 2020). Importantly, in cells infected with CoV2-mut especially at 4 hpi, viral gene translation efficiency relative to cellular genes was reduced (Figure 4b and extended figure 6c). This relative reduction in the translation efficiency of viral genes in cells infected with CoV2-mut virus shows that nsp1, through its binding to the ribosome, promotes viral mRNA translation in infected cells. However, it is important to note that in absolute terms, in cells infected with CoV2-wt the overall translation levels are reduced by approximately 4-fold compared to cells infected with CoV2-mut (Figure 4a), and that ribosome profiling-based measurements provide only relative values of translation. Therefore, it is likely that the absolute translation of viral genes in cells infected with CoV2-mut was comparable or even more efficient than the translation of viral genes in cells infected with CoV2-wt.

### Nsp1 mediates cellular RNA degradation during infection

We next examined the changes in cellular and viral mRNA levels during infection with CoV2-wt and CoV2-mut. To estimate global mRNA levels, we quantified the levels of rRNA and of total RNA extracted from uninfected Calu3 cells and from cells infected with CoV2-wt or CoV2-mut. We observed no substantial differences in either total RNA levels or in rRNA levels (extended data 7a-d). We then sequenced total intracellular RNAs, without rRNA depletion, to assess the relative abundance of cellular and viral mRNAs in uninfected Calu3 and cells infected with the two viruses at 4 and 7 hpi. We found that during infection with CoV2-mut, viral transcripts are produced to the same levels as CoV2-wt (Figure 5a). This shows that even in infected Calu3 cells, during the initial infection, there is no major defect in viral gene expression. However, infection with CoV2-wt was associated with a ~2-fold reduction in cellular mRNA which was not observed in CoV2-mut infected cells (Figure 5a). These results indicate that during infection with CoV2-wt, there is both production of viral transcripts and a concomitant reduction in the levels of cellular transcripts. Whereas, during infection with CoV2-mut the production of viral RNA is maintained but there seems to be no major interference in cellular RNA expression and the levels of cellular mRNAs resemble the levels in mock-infected cells.

**Figure 5:**
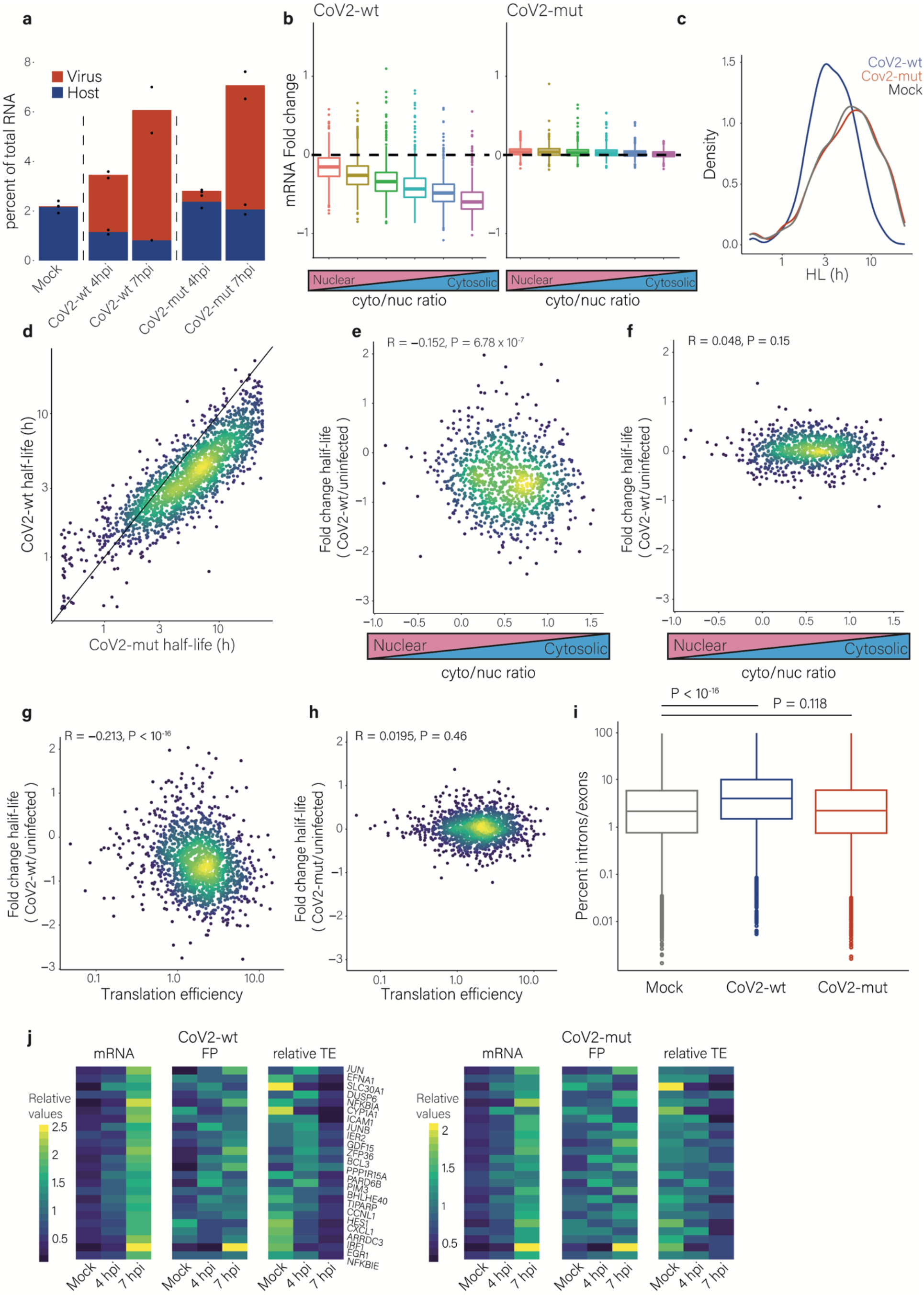
Nsp1 mediates cellular RNA degradation during SARS-CoV-2 infection. **a**, Percentage of reads that aligned to the human or viral transcripts out of total RNA reads. **b**, The fold change in RNA levels in CoV2-wt-infected (left) or CoV2-mut-infected (right) cells relative to uninfected cells at 7 hpi. Transcripts were grouped into six bins on the basis of their cytosol-to-nucleus localization ratio. **c**, The distribution of cellular transcript half-lives in uninfected, CoV2-wt-infected or CoV2-mut-infected Calu3 cells as was determined by SLAM-seq followed by GRAND-SLAM analysis (Jürges et al., 2018). **d**, Scatter plot of cellular transcript half-lives in CoV2-wt-infected cells relative to CoV2-mut-infected cells. (**e** and **f)**, Scatter plots of the fold change of transcript half-lives between CoV2-wt-infected cells (**e**) or CoV2-mut-infected cells (**f**) and uninfected cells, relative to the cytosolic to nuclear ratio in uninfected cells. Pearson’s *R* and two-sided *P* value on log values are presented. (**g** and **h**) Scatter plots of the fold change in transcript half-lives between CoV2-wt-infected cells (**g**) or CoV2-mut-infected cells **(h)** relative to the previously calculated TE values in uninfected cells. Pearson’s *R* and two-sided *P* value on log values are presented. (**i)**, The ratios of intronic to exonic reads for cellular genes in uninfected, CoV2-wt-infected, or CoV2-mut-infected Calu3 cells, at 4 hpi. (**j**) Heatmaps showing the relative mRNA, footprints and translation efficiency, at 4 and 7 hpi of cellular genes that were elevated during infection in both CoV2-wt infected cells and CoV2-mut infected cells relative to uninfected cells.

Since nsp1 mediated mRNA degradation occurs in the cytosol, we examined if the reduction in cellular RNA is associated with their subcellular localization. We found that compared to infection with CoV2-mut, infection with CoV2-wt led to stronger reduction of transcripts that mostly localize to the cytoplasm (*p* value = 5.2×10^−318^ Figure 5b), indicating the reduction in cellular transcript is driven by nsp1-induced degradation that occurs in the cytosol. To directly measure the role of nsp1-ribosome binding to induction of mRNA decay in infected cells, we applied SLAM-seq on uninfected cells and cells infected with CoV2-wt or CoV2-mut. We obtained all characteristics of high-quality SLAM-seq libraries (extended data 8a and 8b), and strong correlation with our previous measurements of mRNA half-life from uninfected and infected cells(Finkel et al., 2021a)(extended data 8c and 8d). Importantly, we observed a substantial reduction in cellular mRNA half-lives upon infection with CoV2-wt, whereas the mRNA half-life in CoV2-mut-infected cells were comparable to the levels measured in mock infected cells (Figure 5c, 5d and extended figure 8e), illustrating that mRNA degradation in infected cells is mediated by nsp1 binding to the ribosome. Furthermore, in agreement with our overexpression experiments, the reduction in half-life correlated with cytosolic localization (Figures 5e and 5f) and with the translation efficiency (Figures 5g and 5h) of host transcripts in cells infected with CoV2-wt, but not in those infected with CoV2-mut. Together these findings demonstrate that the degradation of cellular mRNA during SARS-CoV-2 infection is entirely mediated by nsp1 and is dependent on its interaction with the ribosome.

We previously noticed that SARS-CoV-2 infection led to increased levels of intronic reads relative to exonic reads. However, much of this relative increase may be explained by massive degradation of mature cytosolic mRNAs(Finkel et al., 2021a) and it remains unclear if SARS-CoV-2 directly inhibits splicing as was recently suggested(Banerjee et al., 2020). We therefore examined our data to assess whether increased intronic reads are still detected in cells infected with CoV2-mut, when there is no substantial RNA degradation. Whereas infection with CoV2-wt led to a relative increase in intronic reads, infection with CoV2-mut did not (Figure 5i). These results imply that the increase in intronic reads relative to exonic reads during SARS-CoV-2 infection is largely driven by accelerated degradation of mature cytosolic cellular transcripts by nsp1.

Finally, we previously demonstrated that genes whose mRNA level increase in response to infection did not show a corresponding increase in footprints and we suggested this inability of newly generated transcripts to engage with ribosomes is driven by inhibition of nuclear mRNA export (Finkel et al., 2021a). Inhibition of nuclear mRNA export by SARS-CoV-2 was demonstrated by additional studies (Addetia et al., 2021; Burke et al., 2021; Zhang et al., 2021) and was suggested to be mediated by ORF6 (Addetia et al., 2021) and by nsp1(Zhang et al., 2021), the latter being supported by our overexpression experiments (Figure 2a). Analysis of our mRNA and footprint measurements from infected cells revealed the same signature; mRNAs that were induced in response to infection, did not show a corresponding increase in their footprints. Notably, this occurred with infection with both CoV2-wt and CoV2-mut (Figure 5j), suggesting that the inhibition of nuclear mRNA export indeed represents a distinct function that is independent of nsp1 mediated mRNA degradation and translation inhibition, and is independent on ribosome binding.

Altogether our findings illustrate that nsp1, through induction of mRNA degradation and inhibition of translation, drives host shutoff during SARS-CoV-2 infection. Although nsp1 functional importance lies in blocking the IFN response, its degradation and translation inhibition activities are broad and affect cytosolic mRNAs that are being translated.

### Nsp1 plays a critical role in SARS-CoV-2 pathogenesis *in-vivo*

Viral proteins that suppress host gene expression often play critical roles in viral pathogenesis.To examine the role of nsp1 in SARS-CoV-2 pathogenesis, we utilized the Syrian hamster model, which is susceptible to SARS-CoV-2 infection and exhibits symptoms which are comparable to those seen in humans (Francis et al., 2021; Imai et al., 2020; Rosenke et al., 2020; Sia et al., 2020; Song et al., 2020). We infected two groups of eight hamsters, with CoV2-wt or with CoV2-mut. We measured the weight of the hamsters daily, and collected lung and nasal turbinate (NT) tissues at 3- and 7-days post infection (dpi). Hamsters infected with CoV2-mut did not exhibit a significant change in body weight compared to uninfected hamsters, while CoV2-wt infected hamsters began exhibiting a gradual decrease in body weight from day 2 reaching a statistically significant difference compared to CoV2-mut hamsters at day 6 and remained significant at day 7 (*p* value = 0.0093 and *p* value = 0.0049 by *t-test*, Figure 6a).

**Figure 6:**
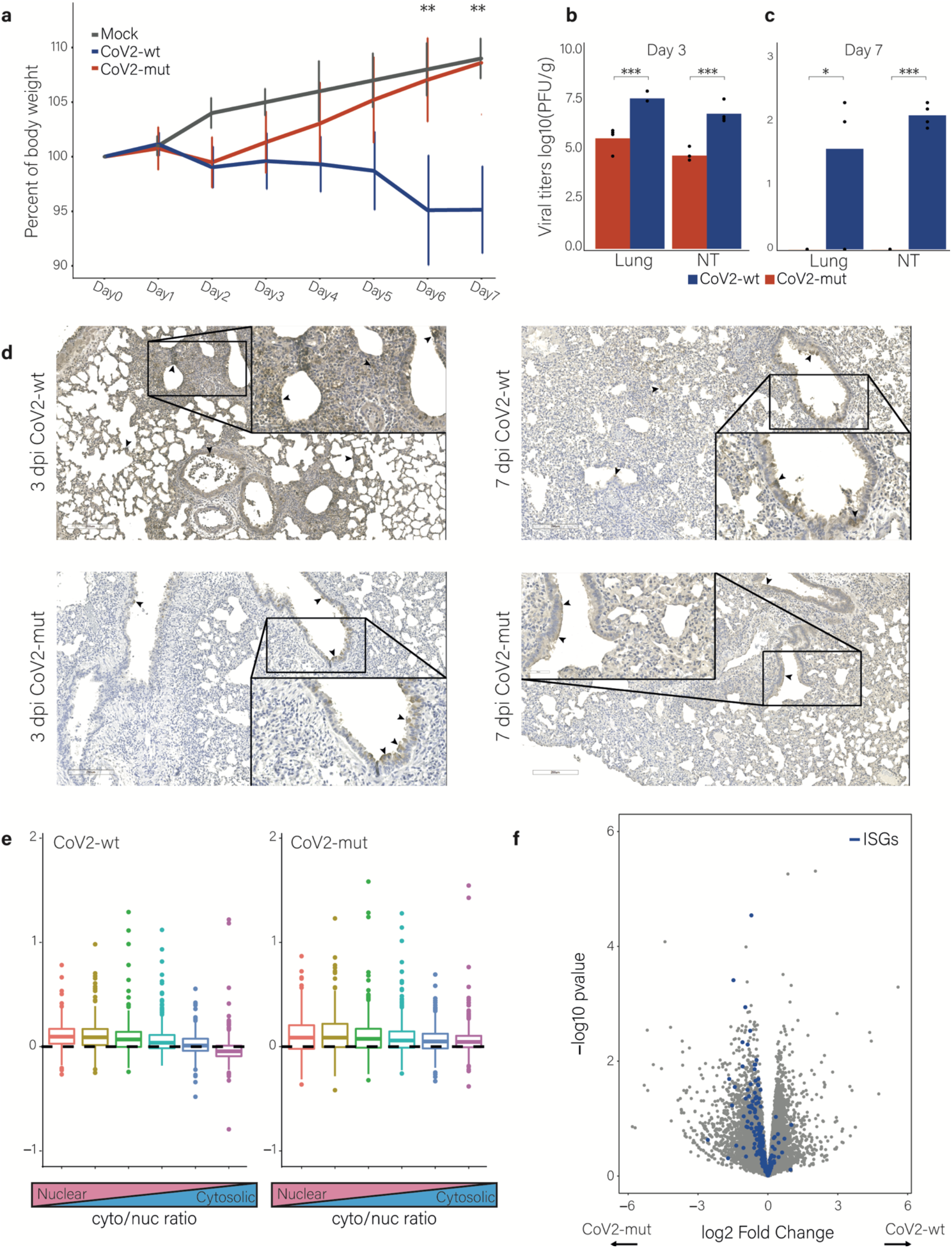
Nsp1 plays a critical role in SARS-CoV-2 pathogenesis in-vivo. **a**, Hamster weight as a percentage of their body weight on day 0 for hamsters infected with CoV2-mut, CoV2-wt, and Mock infected hamsters. (**b** and **c**), Viral titers in lungs and in NT at 3dpi (**b**) and 7dpi (**c**) for both CoV2-wt and CoV2-mut infected hamsters. (**a-c**)Two tailed T-test performed where * = *p*<0.05 and *** = *p*<0.001. **d**, IHC staining of hamster lungs for SARS-CoV-2 spike protein (brown) with hematoxylin counterstain (blue) for hamsters infected with CoV2-wt or CoV2-mut at 3 dpi and 7 dpi. Arrows indicate select infected cells. **e**, The fold change in mRNA levels in CoV2-wt-infected (left) or CoV2-mut-infected (right) relative to mock-infected hamsters at 3 dpi. Transcripts were grouped into six bins on the basis of their cytosol-to-nucleus localization ratio. **f**, Volcano plot showing changes in cellular transcript levels in CoV2-wt versus CoV2-mut infected hamsters at 3 dpi. The fold change and the statistical significance are presented in the x axis and y axis, respectively. ISGs enrichment was calculated using a hypergeometric test. ISGs are marked in blue. two extreme genes were removed.

We next quantified viral titers in lungs and NT in hamsters from both groups. In agreement with the differences in weight loss, at 3 dpi, viral titers were significantly lower in the CoV2-mut-infected hamsters compared to the CoV2-wt-infected hamsters in both the lung (*p* value = 0.0008, t-test) and NT (*p* value = 0.0004, t-test) (Figure 6b). At 7 dpi, CoV2-mut-infected tissues were already cleared from the virus and viral titers were below the threshold of detection, whereas CoV2-wt virus was still significantly detected in both the lung (*p* = 0.0249, t-test) and the NT (*p* < 0.0001, t-test, Figure 6c).

The lungs of infected hamsters were also analyzed by Immunohistochemistry (IHC) staining for SARS-CoV-2 spike protein (Figure 6d). At 3 dpi, CoV2-mut infections were mostly restricted to the epithelial cells of the bronchioles, with minimal staining within the alveoli. In contrast, CoV2-wt infections were not restricted, extending from the bronchioles to the alveoli. In agreement with viral titers measurements, all specimens had less viral staining at 7 dpi than at 3 dpi, indicating resolution of viral infection. While viral infection remained detectable at 7 dpi within the lungs of CoV2-wt challenged hamsters, the infection was largely restricted to the bronchioles. The CoV2-mut 7 dpi specimens appear to have almost recovered from infection, while virus antigen was still detectable it was restricted to the apical surface of the epithelia. We also used RNA-seq to analyze the differences in viral propagation and immune response in the lung at 3 dpi. Lungs from CoV2-mut-infected hamsters exhibited a ~3-fold decrease in the percentage of viral RNA out of the entire mRNA pool when compared to CoV2-wt infected lungs (0.22% compared to 0.69%), further supporting that CoV2-mut propagation, *in-vivo*, is inhibited. We next examined if evidence for nsp1 mediated mRNA degradation could be detected *in-vivo*. Since our results show nsp1-mediated-degradation occurs in the cytosol, we binned hamster transcripts according to the cyto/nuc ratio of their human homologues. We found that in lungs infected with CoV2-wt, and less with CoV2-mut, the levels of transcripts that mostly localize to the cytoplasm (based on homology) were more reduced compared to transcripts that are more nuclear (*p* value = 6.4×10^−27^, Figure 6e), strongly pointing that nsp1-mediated degradation also plays a major role *in-vivo*.

Finally, differential expression analysis revealed a stronger upregulation of ISGs in the lungs of CoV2-mut infected animals compared to CoV2-wt-infected lungs (*p* value = 0.009, Figure 6f). This indicates that in accordance with our cell culture findings, animals infected with nsp1 mutant virus induced a stronger type I interferon response compared to animals infected with the WT virus.

Overall, these results show that nsp1, through its binding to the ribosome, plays a critical role in the SARS-CoV-2 life cycle by interfering with host innate immune responses, supporting efficient propagation in the lungs and ensuing disease.

## Discussion

Many viruses have developed varied and sophisticated mechanisms to repress host mRNA translation while concomitantly allowing the translation of viral mRNAs. In the case of SARS-CoV-2, several viral ORFs have been suggested to interfere with cellular gene expression, including nsp1. We show here that nsp1 is responsible for the shutoff in protein synthesis during SARS-CoV-2 infection, and its activity is mediated by three distinct mechanisms: translation block, induction of mRNA degradation and interference in nuclear mRNA export. Using reporters and broad measurements of translation and degradation promoted by WT and nsp1-mutant viruses, we show that the degradation and translation inhibition activities occur in the context of an nsp1-ribosome interaction whereas the nuclear export activity does not significantly contribute to the global shutoff observed in infected cells, and is independent from nsp1 binding to the ribosome.

Despite numerous studies, it remains unclear how viral transcripts overcome nsp1-mediated translation suppression and specifically how this evasion relates to the translation and degradation activities of nsp1. Our results together with recent studies (Bujanic et al., 2021; Finkel et al., 2021a; Mendez et al., 2021; Vora et al., 2022) show that in the setting of nsp1 overexpression, cellular mRNA translation is strongly inhibited by nsp1. The observation that nsp1-CD, which does not lead to mRNA degradation, inhibits translation to a lesser extent than WT nsp1, suggests that both the translation inhibition and the degradation activities contribute to nsp1’s ability to shutoff cellular protein synthesis. It is clear that the viral 5’leader sequence, which is shared by all viral transcripts, allows transcripts to evade the inhibition imposed by nsp1 overexpression, but whether this reflects evasion from nsp1 induced degradation, evasion from the nsp1-mediated block in translation or evasion from both activities is harder to dissect. Our measurements, in combination with recent work (Finkel et al., 2021a; Mendez et al., 2021), clearly show that the viral 5’ leader sequence makes mRNAs refractory to nsp1-mediated degradation. With regard to evasion from nsp1 translational block, two models could potentially explain viral mRNA escape. One posits that nsp1 blocks the translation of both cellular and viral transcripts, but since cellular transcripts are degraded, viral mRNAs remain the only available target for ribosomes that are not inhibited by nsp1. The other model proposes that viral mRNAs are refractory to suppression by nsp1, as they interact with nsp1 in a manner that causes a change in its conformation, removing its C-terminal domain from the ribosome entry channel and facilitating viral RNA translation (Mendez et al., 2021; Shi et al., 2020; Tidu et al., 2020). These two models may not be mutually exclusive and both mechanisms may contribute to the evasion of viral mRNAs from the nsp1 translational block. The observation that nsp1-CD, which contains two point mutations in the N-terminal region, does not degrade cellular mRNAs, and in contrast to nsp1-WT also inhibits the translation of mRNA containing the viral 5’ leader sequence, could be reconciled by both of these models. If these specific mutations in the N-terminal region affect the ability of nsp1 to interact and recognize the viral 5’ leader sequence as was recently suggested (Mendez et al., 2021), it would explain why translation of viral mRNAs is inhibited by this nsp1 mutant. On the other hand, since nsp1-CD does not lead to cellular mRNA degradation, the competition of viral mRNAs over ribosomes that are not blocked by nsp1 will grow, and this can also contribute to reduction in viral mRNA translation. Our measurements from CoV2-wt-infected cells indicate that viral mRNAs are not preferentially translated compared to cellular transcripts. However, a major limitation of these measurements is that viral mRNA replication and transcription is massive, and occurs within a unique endomembrane system (Wolff et al., 2020) that would thus render much of the viral mRNA inaccessible to ribosomes. This inaccessibility of potentially large portions of viral RNA will skew our measurements and it is likely the true translation efficiency of viral transcripts is significantly higher. Comparing the translation efficiencies of viral and cellular transcripts in cells infected with CoV2-wt or with CoV2-mut allowed us to overcome this limitation and to demonstrate that in infected cells nsp1 indeed specifically promotes the translation of viral transcripts compared to cellular transcripts, as we observed that in cells infected with CoV2-mut, viral genes are translated even less efficiently compared to their cellular counterparts. However, at the same time our measurements of the overall translation capacity in infected cells, using incorporation of OPP into nascent chains, show that nsp1-WT leads to a major reduction in the absolute levels of translation in infected cells. Since the vast majority of mRNA in infected cells is viral mRNA this means that viral mRNAs translation is likely also inhibited in the presence of nsp1. Together these results support a model in which nsp1 acts as a strong inhibitor of translation that tightly binds to ribosomes and thus inhibits the translation of both viral and cellular mRNAs. Viral transcripts through their 5’leader can at least partially escape this translation repression, making them less sensitive to nsp1 mediated translation inhibition. At the same time, ribosome bound nsp1 also leads to accelerated degradation of cellular but not of viral mRNAs providing means for viral mRNAs to quickly take over and dominate the mRNA pool. The accumulation of SARS-CoV-2 mRNAs and the ability of these transcripts to partially evade nsp1 mediated repression explains how infected cells so effectively shutoff the synthesis of cellular proteins while switching to translation of viral mRNAs.

We reveal here that similarly to SARS-CoV nsp1 (Kamitani et al., 2009; Lokugamage et al., 2012), SARS-CoV-2 nsp1 likely induces endonucleolytic cleavage of cellular mRNA that leads to their degradation, and this degradation activity depends on nsp1 binding to ribosomes. Our broad measurements, both by over-expression and in the context of infection, show that nsp1 accelerates the degradation of cytosolic RNAs, and specifically the ones that are more highly translated. Since nsp1 is physically bound to the ribosome and the degradation depends on this interaction, an appealing speculation is that nsp1 recruits a cellular nuclease whose activity relates to mRNA translation surveillance. A major future challenge will be to identify this nuclease.

Host shutoff is traditionally thought to contribute to viral propagation by facilitating viral protein translation and by blocking the ability of the cell to mount an efficient antiviral response. Our results illustrate that the main functional importance of host shutoff during SARS-CoV-2 infection is the latter. The observation that the CoV2-mut virus doesn’t have a substantial propagation defect in Vero cells, which lack the ability to produce IFN, together with our microscopy, RNA-seq and Western blot analysis showing equivalent accumulation of viral proteins, illustrates that viral gene expression is intact in cells infected with CoV2-mut. This intact gene expression strongly suggests that the growth defect of CoV2-mut in Calu3 cells and *in-vivo* in infected hamsters, is not due to a direct reduction in viral gene translation but rather due to the absence of shutoff of cellular protein synthesis.

Furthermore, analysis of the early time points in infected Calu3 cells, which have an intact IFN response, also illustrates that in the absence of a functional nsp1 there is no major block in the initial viral protein production during the first infection cycle. These results suggest that the defects observed in CoV2-mut propagation are due to the critical role nsp1 plays in blocking type I IFN response, which will affect neighboring cells and the ability of the virus to spread. These observations also coincide with the observed attenuation and inhibition in spread of CoV2-mut virus in infected hamsters. Activation of the type I IFN response leads to the induction of hundreds of ISGs in infected as well as in neighboring cells, directing cells into an antiviral state. It is therefore likely that the stronger induction of ISGs during infection with CoV2-mut represents elevated levels of type I IFN production. Since SARS-CoV-2 is sensitive to IFN treatment (Felgenhauer et al., 2020; Katsura et al., 2020; Mantlo et al., 2020; Shemesh et al., 2021; Vanderheiden et al., 2020), differences in IFN production, which affects both infected cells and neighboring cells, can explain the inhibition in CoV2-mut propagation. Supporting this notion are recent findings showing nsp1-deficient replicons are more sensitive to IFN treatment (Ricardo-Lax et al., 2021).

Deletion of nsp1-coding sequence in MHV, revealed nsp1 is a major virulence factor in mice and it plays a critical role in blocking the IFN response (Shen et al., 2019; Züst et al., 2007). SARS-CoV nsp1 was also shown to be a pathogenic determinant and to play a role in the inhibition of host antiviral response in cell culture(Jimenez-Guardeño et al., 2015; Wathelet et al., 2007). These findings together with our results illustrate that nsp1 acts as a major virulence factor in CoVs. Since the loss of nsp1 function renders the virus vulnerable toward immune clearance, it may serve as an attractive therapeutic target that will be effective against many CoVs and SARS-CoV-2 variants.

Overall our data establish that nsp1 is a major immune evasion factor of SARS-CoV-2. Although SARS-CoV-2 encodes additional inhibitors of the innate immune defenses, and specifically the IFN response (Xia et al., 2020), we show that a loss in nsp1 activity renders the virus vulnerable to immune response both *in-vitro* in cells with intact IFN response and *in-vivo* in hamsters. Mechanistically we illustrate that in the context of infection, nsp1 mediates its function by both induction of cellular mRNA degradation and inhibition of translation, these functions are linked and it is likely that both are major contributors to SARS-CoV-2 virulence. In conclusion, our study provides an in-depth picture of how nsp1 interferes with cellular gene expression, and reveals the critical role it plays in SARS-CoV-2 pathogenesis specifically through shutoff of the IFN response.

## Acknowledgments

We thank Igor Ulitsky and Schraga Schwartz for providing valuable feedback. This work was supported by Public Health Service grant AI146081 from the National Institutes of Health and pilot grants from the Institute for Human Infections and Immunity at The University of Texas Medical Branch to SM; By the National Institutes of Allergy and Infectious Diseases Center for Research on Influenza Pathogenesis (HHSN272201400008C), Research Program on Emerging and Re-emerging Infectious Diseases (JP20fk0108412) and the Japan Program for Infectious Diseases Research and Infrastructure (JP21wm0125002) from the Japan Agency for Medical Research and Development (AMED) to YK; By research grants from the Weizmann Corona Response Fund, the Knell Family Center for Microbiology and by a research grant from the Weizmann SABRA - Yeda-Sela - WRC Program, the Estate of Emile Mimran, and The Maurice and Vivienne Wohl Biology Endowment to NS-G.

## Supplemental figures

**Extended data figure 1:**
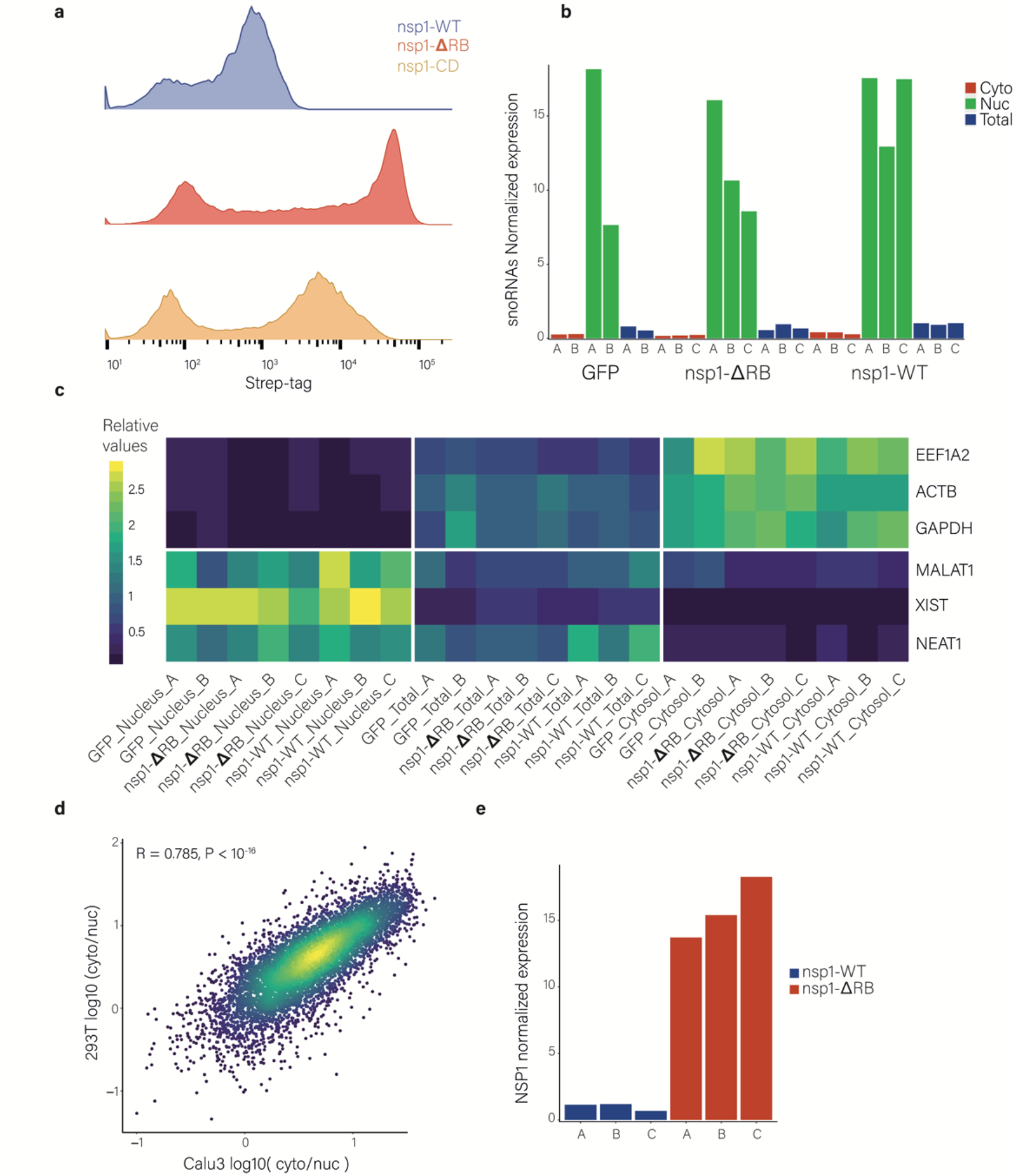
Nsp1 inhibits its own expression in transfected cells. **a**. Flow cytometry analysis of the Strep tag expression in cells transfected with nsp1-WT, nsp1-ΔRB, or nsp1-CD containing strep-tag at the C-terminus. **b**. Normalized sum of the expression of snoRNAs in the nuclear, cytosolic, and total fractions of 293T cells transfected with nsp1-WT, nsp1-ΔRB, or GFP. **c**. Heatmap showing the relative expression of cellular mRNAs and nuclear lncRNAs in the nuclear, cytosolic, and total fractions of 293T cells transfected with nsp1-WT, nsp1-ΔRB, or GFP. **d**. Scatter plot of transcript cyto/nuc ratios calculated from fractionation measurements in 293T cells transfected with plasmid expressing GFP relative to cyto/nuc ratios measured in Calu3 cells (Finkel et al., 2021a). R (pearson) and two-sided P value are presented. **e**. Expression of transcripts encoding nsp1 calculated from RNA-seq data. Expression of nsp1-WT transcripts or nsp1-ΔRB transcripts was normalized to NEAT1 RNA (a nuclear long non coding RNA), and nsp1-ΔRB expression relative to the mean of nsp1-WT samples are presented.

**Extended data figure 2:**
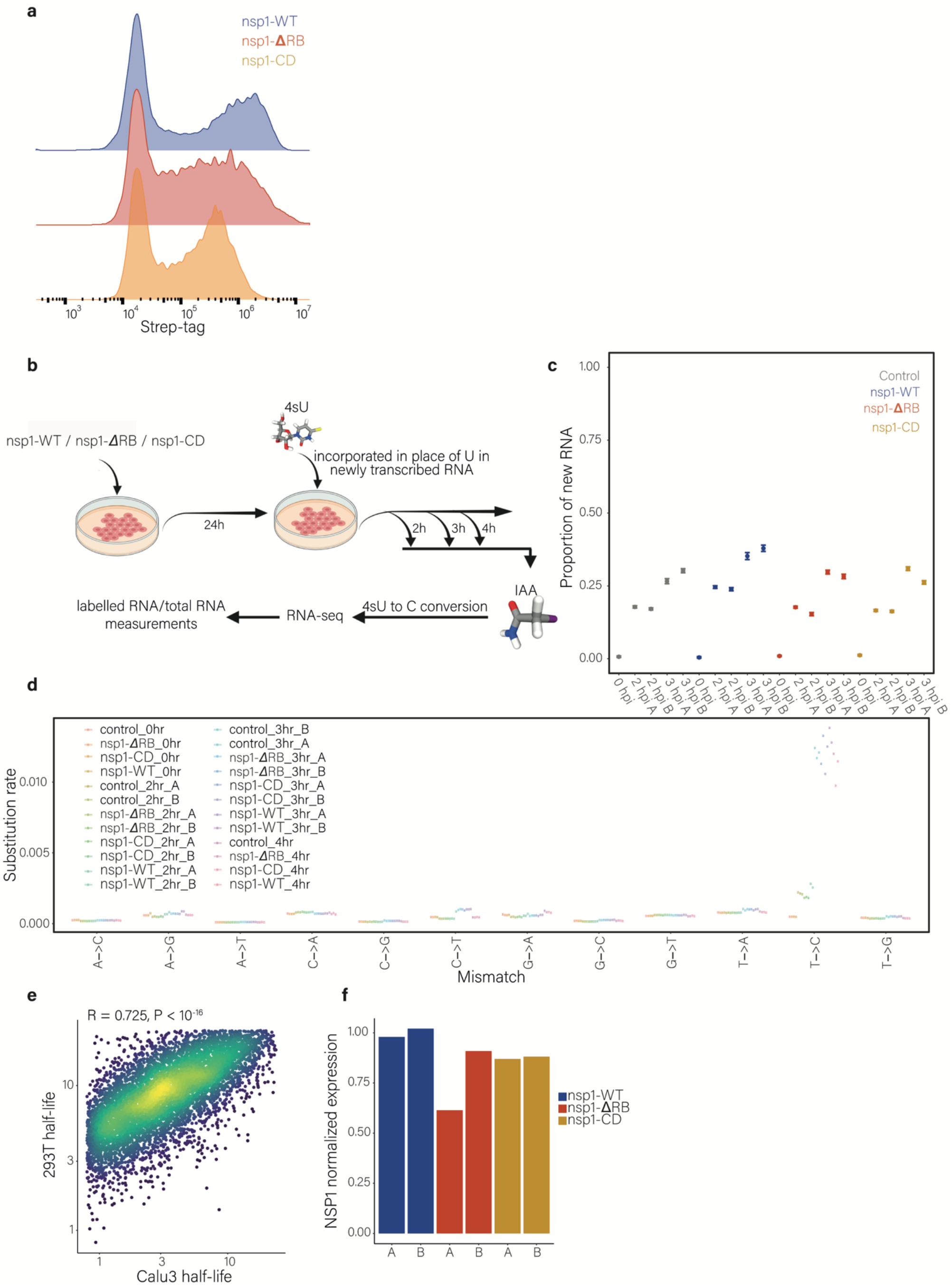
Subcellular fractionation and SLAM-seq measurements in nsp1 expressing cells. **a**. Flow cytometry analysis of the Strep tag-positive cells, in cells transfected with plasmid expressing nsp1-WT, nsp1-ΔRB, or nsp1-CD, all of which were cloned downstream to the CoV2 viral 5’ leader sequence. **b**. Experimental design of the SLAM-seq measurements and the half-life calculation. **c**. Rates of nucleotide substitutions demonstrate efficient conversion rates in 4sU-treated samples compared to non-treated cells (no 4sU). **d**. Proportion of new-to-total RNA (NTR) in 293T cells transfected with plasmid expressing GFP, nsp1-WT, nsp1-ΔRB, or nsp1-CD as calculated by GRAND-SLAM(Jürges et al., 2018). **e**. Scatter plot of transcript half-lives calculated from SLAM-seq measurements in 293T cells transfected with plasmid expressing GFP relative to transcript half-lives in Calu3 cells(Finkel et al., 2021a). Pearson’s R and two-sided P-value are presented. **f**. Expression levels of nsp1 transcripts calculated from RNA-seq data. Nsp1 expression in each of the conditions (nsp1-WT, nsp1-ΔRB, or nsp1-CD) was normalized to NEAT1 expression, and nsp1 expression relative to the mean of nsp1-WT samples are presented.

**Extended data figure 3:**
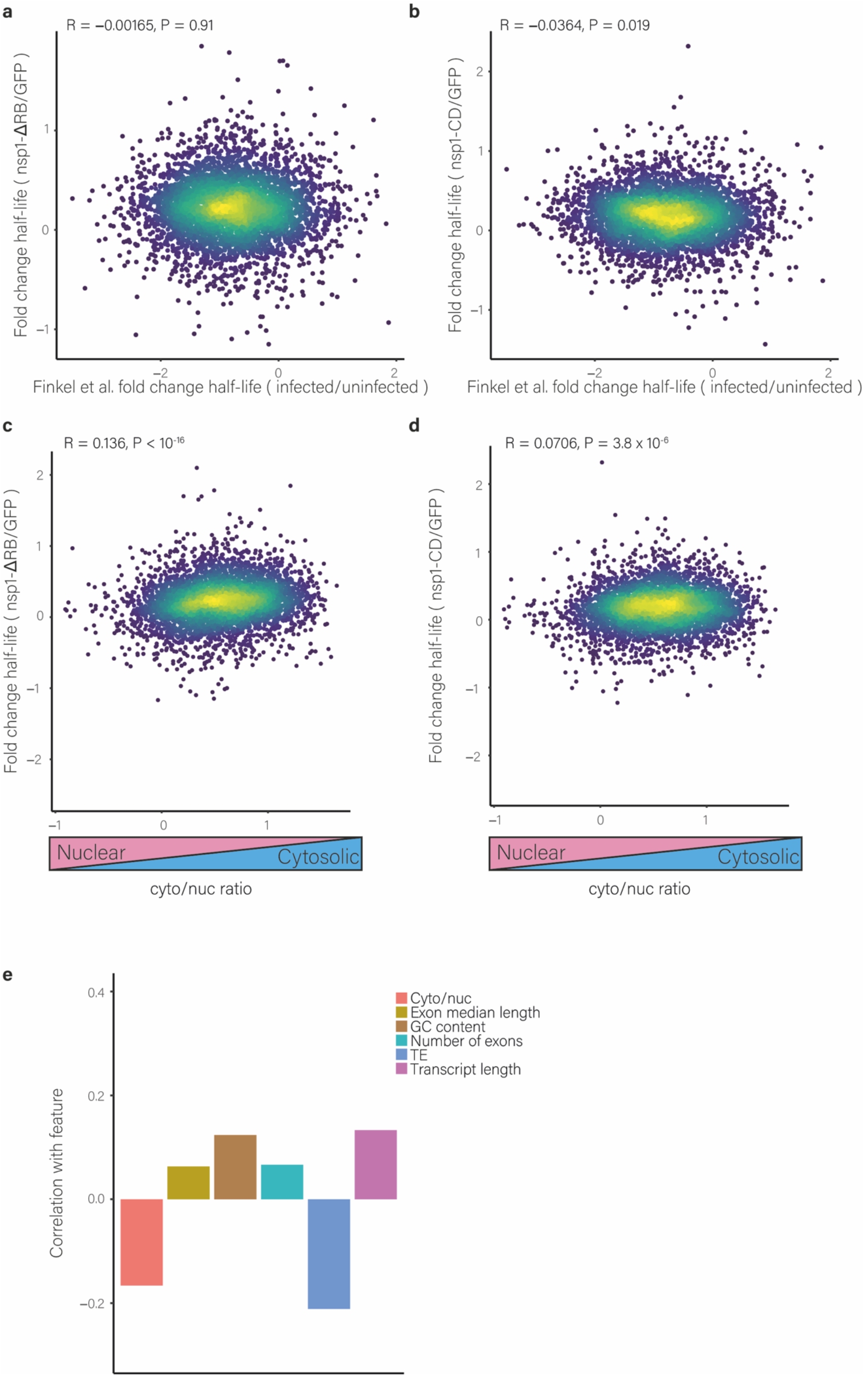
Analysis of gene expression in cells expressing nsp1-ΔRB or nsp1-CD. **a and b**. Scatter plot of fold change in transcript half-lives in 293T cells transfected with plasmid expressing nsp1-ΔRB (**a**) or nsp1-CD (**b**) relative to those transfected with plasmid expressing GFP and the fold change transcript half-lives in uninfected and infected cells(Finkel et al., 2021a). Pearson’s R and two-sided P values are presented. **c and d**. Scatter plot of fold change in transcript half-lives in 293T cells transfected with a plasmid expressing nsp1-ΔRB (**c**) or nsp1-CD (**d**) relative to those transfected with a plasmid expressing GFP and cyto/nuc ratios of uninfected cells (Finkel et al., 2021a). Pearson’s R and two-sided P value are presented. **e**. Correlation between gene features and the fold change of transcript half-lives between 293T transfected with a plasmid expressing nsp1-WT or control plasmid.

**Extended data figure 4:**
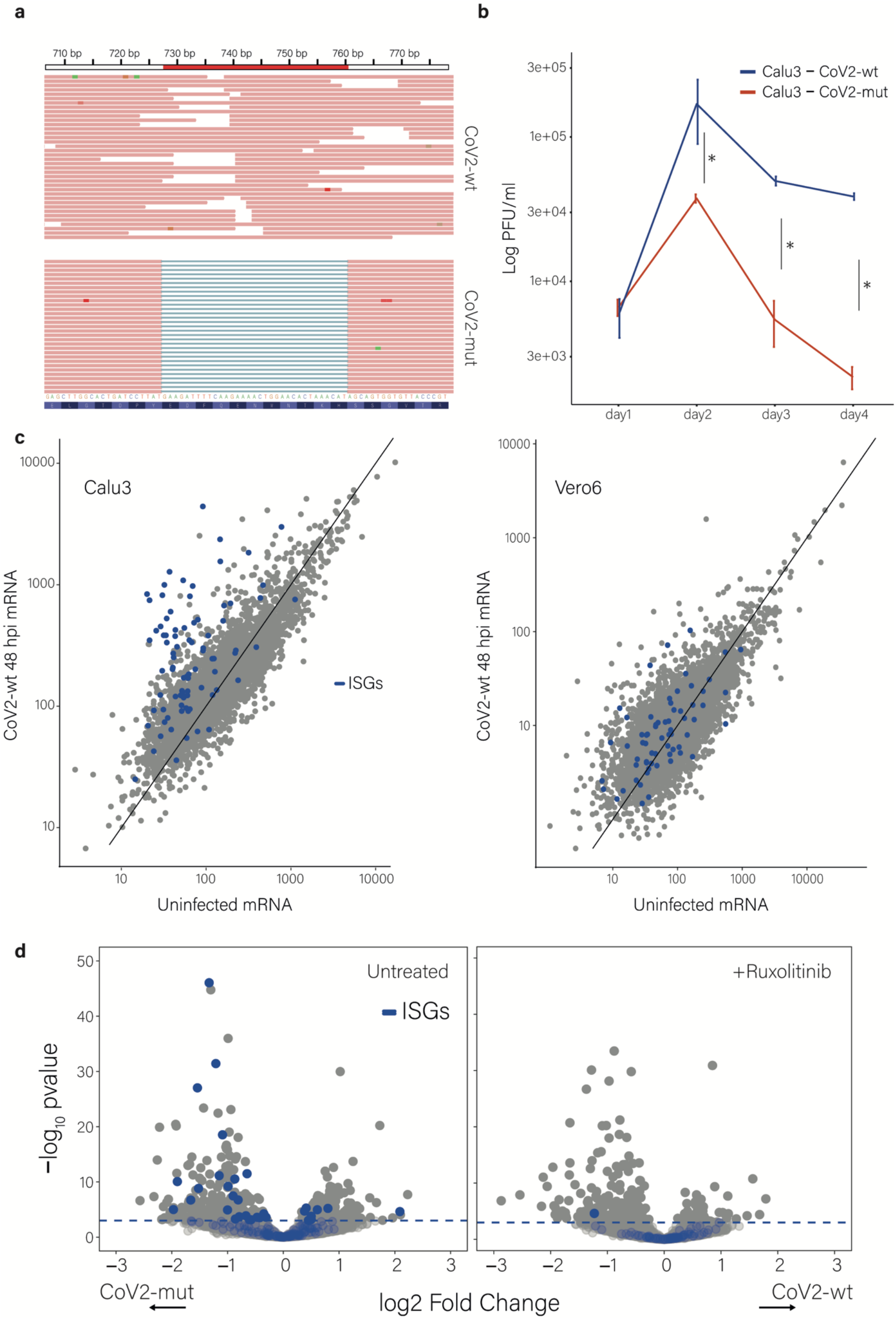
Gene expression and virus growth kinetics in CoV2-wt-infected cells and CoV2-mut-infected cells. **a**, RNA reads spanning the deletion region in nsp1 (nt 728-760) in the SARS-CoV-2-WT (CoV2-wt) or in SARS-CoV-2 carrying nsp1-ΔRB mutant (CoV2-mut). **b**. Viral titers in supernatant collected from Calu3 cells infected with CoV2-wt or with CoV2-mut (MOI=0.01) at 1, 2, 3, and 4 dpi. The titers of the CoV2-wt infection were higher than the titers of the CoV2-mut infection at all timepoints (two tailed t-test, where * = *p*<0.01) **c**. Scatter plot plotting the TPM from uninfected Calu3 or Vero cells and cells infected with CoV2-wt at 48 hpi (MOI = 0.01). ISGs (blue) and all the rest of genes (gray) are marked on each of the plots. **d**. Volcano plots showing changes in cellular transcript levels in CoV2-wt versus CoV2-mut infected Calu3 cells that were left untreated (left) or treated with Ruxolitinib (right) at 48 hpi. The dotted line represents the cutoff for significant genes (p=0.001) and ISGs are marked in blue.

**Extended data figure 5:**
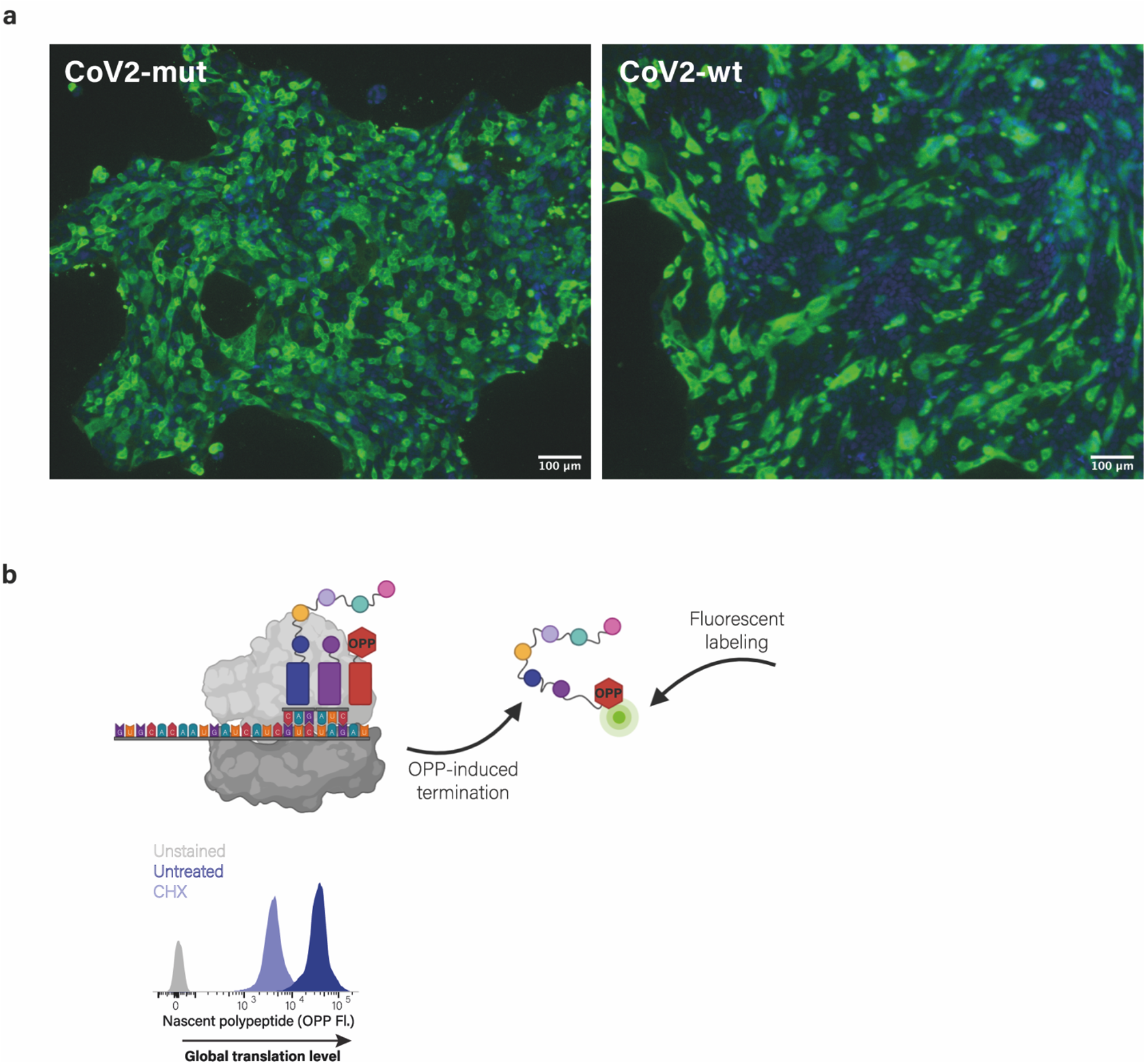
Similar levels of infectivities of CoV2-wt and Cov2-mut in Calu3 cells (a) and measurement of translational activities by using OPP incorporation (b). **a**. Microscopy images of Calu3 cells infected with CoV2-wt or CoV2-mut at an MOI of 3 at 8hpi stained with antisera against SARS-CoV-2 (green) and DAPI (blue). **b**. Scheme of labeling and detection of nascent protein synthesis by OPP incorporation followed by fluorescent labeling. OPP is efficiently incorporated into newly translating proteins, releasing the polypeptides and terminating translation. Following fixation, OPP is fluorescently labeled using a Click reaction and can then be measured by flow cytometry. Flow cytometry analysis of untreated 293T cells and 293T cells treated with cycloheximide leading to efficient translation inhibition. Unstained cells are shown as control.

**Extended data figure 6:**
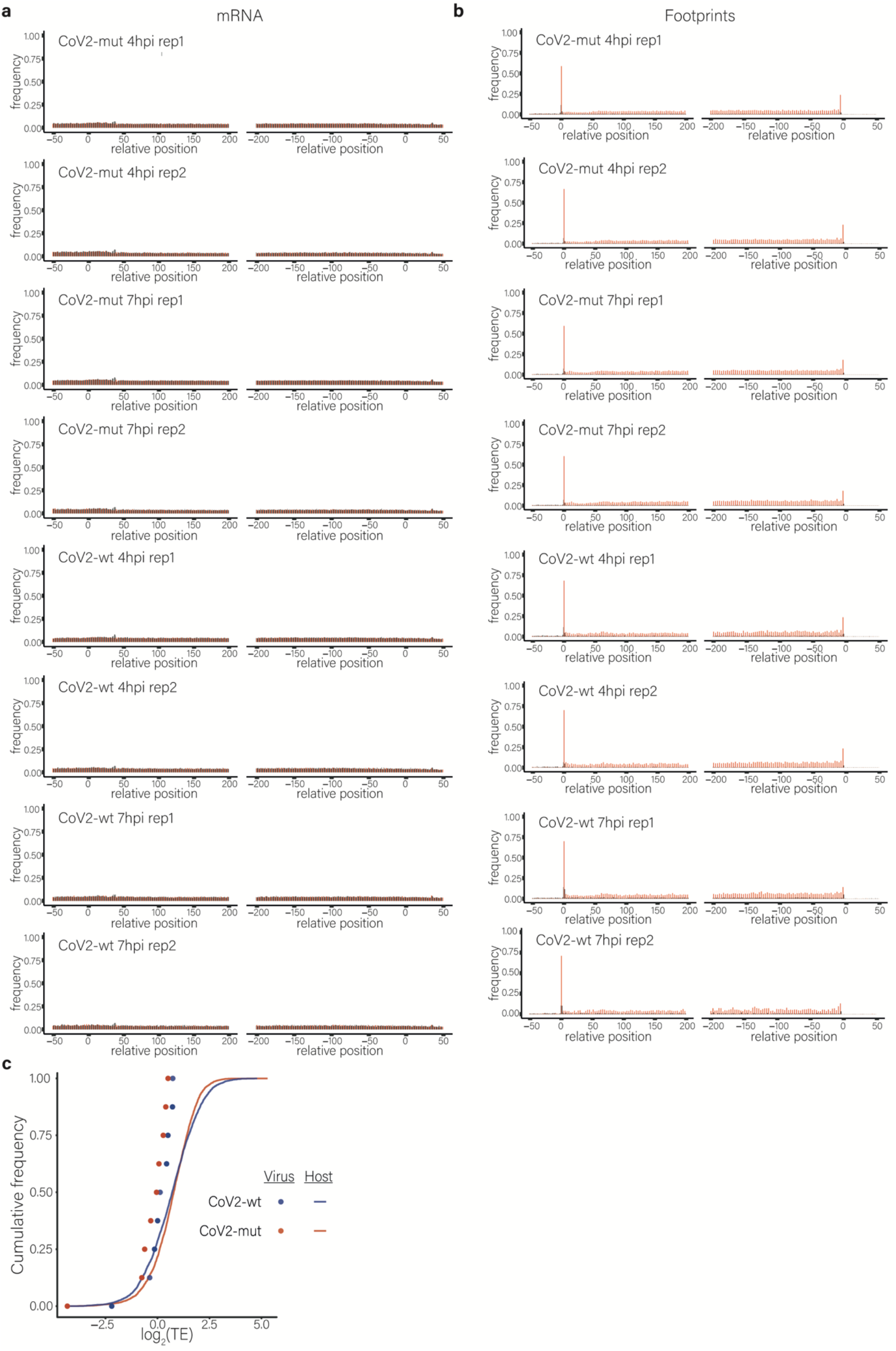
Ribosome profiling measurements in CoV2-wt-infected cells and Cov2-mut-infected cells. **a and b**. Metagene analysis for the mRNA (**a**) and footprint (**b**) samples. **c**. Cumulative frequency of human (line) and viral (dots) genes according to their relative TE in cells infected with CoV2-wt (blue) or with CoV2-mut (red) at 7 hpi. TE was calculated from ribosome profiling in parallel to mRNA sequencing, and is defined as the ratio of ribosome footprints to mRNA for a given gene. Each dot represents one of nine major viral mRNA species.

**Extended data figure 7:**
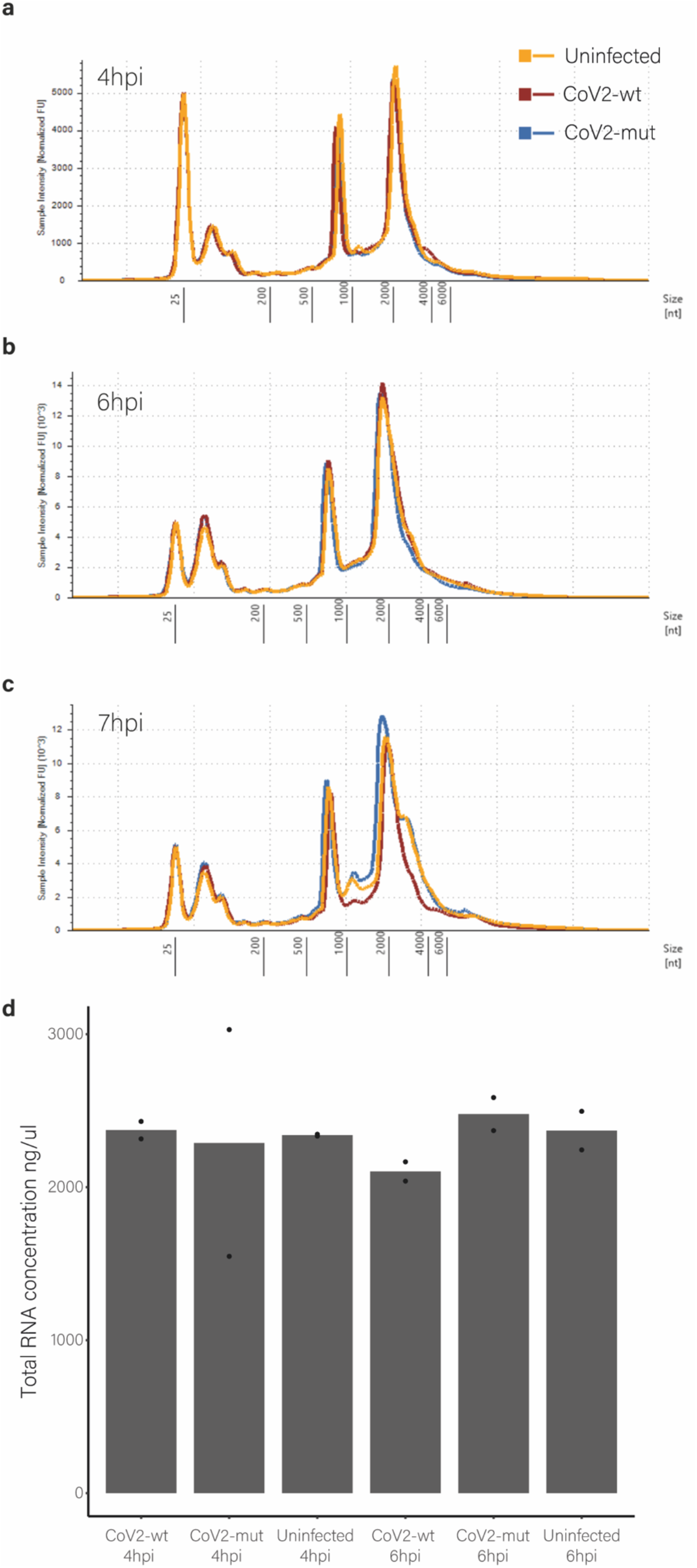
Amounts of RNA in CoV2-wt and CoV2-mut infected cells. **a-c**. The concentration of the 18S and 28S ribosomal RNAs were measured in uninfected Calu3 cells and Calu3 cells infected with CoV2-wt or CoV2-mut at 4 hpi (a), 6 hpi (b), or 7 hpi (c) by using a TapeStation system. **d**. Total RNA was measured in uninfected Calu3 cells and in CoV2-wt or CoV2-mut infected Calu3 cells at 4 and 6 hpi by Qubit Fluorometer.

**Extended data figure 8:**
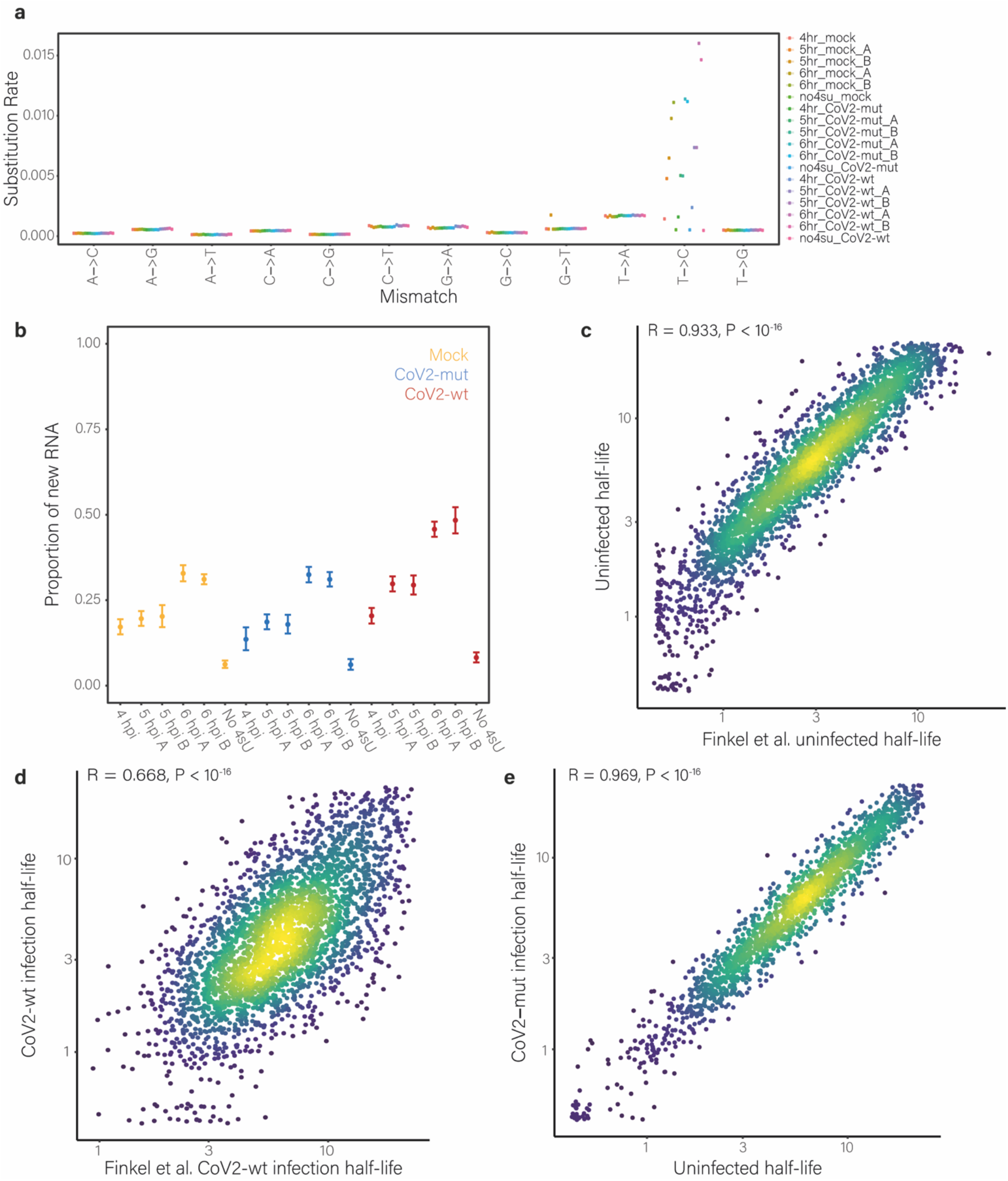
SLAM-seq measurements in CoV2-wt-infected cells and CoV2-mut infected cells. **a**. Rates of nucleotide substitutions demonstrate efficient conversion rates in 4sU-treated samples compared to non-treated cells (no 4sU). **b**. Proportion of new-to-total RNA (NTR) in Calu3 cells infected with CoV2-wt or CoV2-mut as calculated by GRAND-SLAM(Jürges et al., 2018). **c** and **d**. Scatter plot of transcript half-lives in (**c**) uninfected Calu3 and (**d**) CoV2-wt-infected cells relative to our previous half-lives measurements from uninfected and infected cells (Finkel et al., 2021a). Half-lives lower than 0.8 h or higher than 24h were omitted. Pearson’s R and two-sided P value are presented. **e**. Scatter plot of transcript half-lives in CoV2-mut-infected Calu3 cells relative to half-lives in mock infected cells. Pearson’s R and two-sided P value are presented.

## Methods

### Cells and viruses

Calu3 cells (ATCC HTB-55) were cultured in 6-well or 10-cm plates with Dulbecco’s Modified Eagle Medium (DMEM) supplemented with 10% fetal bovine serum (FBS), MEM non-essential amino acids (NEAA), 2 mM L-glutamine, 100 units per ml penicillin and 1% Na-pyruvate. African green monkey kidney clone E6 cells (Vero E6, ATCC® CRL-1586™) were grown in growth medium DMEM containing 10% FBS, MEM NEAA, 2 mM L-glutamine, 100 Units/ml penicillin, 0.1 mg/ml streptomycin, 12.5 Units/ml nystatin (P/S/N), all from Biological Industries, Israel]. Calu3 monolayers were washed once with MEM-Eagles medium (MEM) without FBS and infected with SARS-CoV-2 in the presence of 20 μg per ml TPCK trypsin (Thermo scientific). Plates were incubated for 1 h at 37°C to allow viral adsorption. Then, MEM medium supplemented with 2% FBS, was added to each well. Virus was propagated and titered on Vero E6 cells and sequenced prior to use. Handling and working with SARS-CoV-2 were conducted in a BSL3 facility in accordance with the biosafety guidelines of the Israel Institute for Biological Research (IIBR), or in a BSL3 facility at The University of Texas Medical Branch (UTMB). The Institutional Biosafety Committee of Weizmann Institute and that of UTMB approved the protocol used in these studies. Calu3 and 293T cells were authenticated by ATCC using STR profiling. All cell lines tested negative for mycoplasma.

### Plasmid construction and generation of recombinant infectious SARS-CoV-2

SARS-CoV-2 nsp1 ORF, carrying a C-terminal myc-tag, was cloned into pCAGGS-MCS, resulting in pCAGGS-nsp1-WT. Similarly, SARS-CoV-2 nsp1 carrying a deletion of the amino acids 155 to 165, and SARS-CoV-2 nsp1 carrying R124A and K125A mutations, were cloned into pCAGGS to generate pCAGGS-nsp1-ΔRB and pCAGGS-nsp1-CD, respectively. The plasmid pRL-EMCV-FL, carrying an upstream, cap-dependent Renilla luciferase gene followed by an EMCV IRES-dependent firefly luciferase gene, was constructed, as described previously (Harcourt et al., 2020; Narayanan et al., 2015). To generate CoV-2-mt, carrying a deletion of the amino acids 155 to 165 in the nsp1 gene of SARS-CoV-2 genome, a recombinant PCR procedure was employed to generate the deletion in fragment 1 of the SARS-CoV-2 reverse genetics system (Xie et al., 2020). The SARS-CoV-2 full-length cDNA was assembled from fragments 1 through 7 and used as the template for synthesizing the full-length RNA transcripts as described previously (Xie et al., 2020). The recombinant infectious viruses, CoV-2-wt and CoV-2-mt, were recovered using the full-length RNA transcripts, according to established protocols, as described previously (Xie et al., 2020).

### Virus infection and quantification

Recombinant wild-type SARS-CoV-2 (CoV-2-wt) and mutant SARS-CoV-2 (CoV-2-mt), both of which were rescued by using the reverse genetics system (Xie et al., 2020), were passaged once in Vero E6 cells, and used for infection studies. For virus growth analysis, Vero E6 and Calu3 cells were infected at the indicated m.o.i. for 1 h at 37°C. After virus adsorption, cells were washed twice and incubated with the appropriate medium. Culture supernatants were collected at the indicated times post-infection and the infectious virus titers were determined by plaque assay on Vero E6 cells, as described previously (Harcourt et al., 2020).

### Luciferase assay

Vero E6 cells in 24-well plates were co-transfected, in triplicate, with pRL-EMCV-FL reporter plasmid and the indicated pCAGGS-based nsp1 expression plasmids using TransIT-LT1 reagent (Mirus). At 24 hours post-transfection, the cells were lysed in Passive Lysis Buffer (Promega), and luciferase assays were performed using the Promega Dual Luciferase assay system.

### Viral titers

For determination of growth rate of SARS-CoV-2 WT or SARS-CoV-2 mut viruses, Vero E6 and Calu3 cells were seeded in 12-well plates (5E5 and 1E6 cells/well respectively). 24 hours later Calu3 cells were washed once with MEM with NEAA, glutamine, and P/S/N without FBS. Then, Calu3 and Vero E6 cells were infected with 0.1 MOI of SARS-CoV-2 wt or SARS-CoV-2 mut viruses in infection medium (MEM containing 2% FBS with NEAA, glutamine, and P/S/N). For Calu3 cells, the infection medium was supplemented with 20 μg per ml TPCK trypsin (Thermo scientific). Plates were incubated for 1 h at 37°C to allow viral adsorption. Then, the infection medium was aspirated, the wells were washed once with infection medium and then MEM medium supplemented with 2% FBS and 0.15% Sodium bicarbonate, was added. Culture media was collected at several time points: 0 hours, 11 hours, 24 hours and 48 hours. Cell debris were removed by centrifugation. The supernatants were kept for viral titer determination and the cell pellets were harvested by Tri-reagent (Merck, Israel) for RNA purification.

For viral titration, Vero E6 cells were seeded in 12-well plates (5E5 cells/well) and grown overnight in growth medium. Serial dilutions of each supernatant from each of the above indicated time points were prepared in infection medium and used to infect Vero E6 monolayers (200 μl/well). Plates were incubated for 1 h at 37 °C to allow viral adsorption. Then, 2 ml/well of overlay [MEM containing 2% FBS and 0.4% tragacanth (Merck, Israel)] was added to each well and plates were incubated at 37 °C, 5% CO2 for 72 h. The media were then aspirated, and the cells were fixed and stained with 1 ml/well of crystal violet solution (Biological Industries, Israel). The number of plaques in each well was determined, and SARS-CoV-2 WT and SARS-CoV-2mut titers were calculated.

### Experimental Infection of Syrian Hamsters

All hamster experiments with SARS-CoV-2 were performed under biosafety level-3 agriculture (BSL-3 Ag) containment at the University of Wisconsin-Madison in laboratory space approved by the Centers for Disease Control and Prevention and by the US Department of Agriculture.

Golden Syrian hamsters (females; 4-5 weeks old) were intranasally infected with 1000 pfu of virus in 50 μl of inoculum while under isoflurane anesthesia. Animals were checked daily to evaluate the health of the hamsters, including body weight. Three days and seven days after infection, tissues were collected after the animals were humanely euthanized.

Virus titers in the tissues were determined on confluent Vero E6/TMPRSS2 cells obtained from the Japanese Collection of Research Bioresources (JCRB) Cell Bank (1819). Cells were infected with 100 μl of undiluted or 10-fold dilutions (10^−1^ to 10^−5^) of clarified lung and nasal turbinate homogenates for 30 min. After the inoculum was removed, the cells were washed once, and then overlaid with 1% methylcellulose solution in DMEM with 5% FBS. The plates were incubated for three days, and then the cells were fixed and stained with 20% methanol and crystal violet in order to count the plaques.

### Immunohistochemistry staining for hamster tissues

Virally infected cells within the lungs were detected as described before (Tseng et al., 2007; Yoshikawa et al., 2009). Briefly, paraffin-embedded sections were firstly subjected to immunostaining with a commercially available and diluted (1:5,000) rabbit-anti-SARS-CoV-2 spike (S) protein (ab272504, Abcam Plc., Cambridge, UK), followed by staining with a peroxidase-conjugated secondary antibody. Thoroughly washed tissues were then incubated with a solution containing 3,3’-Diaminobenzidine as the substrates to visualize virally infected cells, according to the procedure recommended by the vendor (MP-7802, Vector Laboratories, Burlingame, CA). Finally, stained slides were counterstained with hematoxylin and antigen expression was examined under an inverted microscopy at different magnifications.

### Immunofluorescence analysis

Vero E6 cells were either uninfected or infected with CoV2-wt or CoV2-mt at an moi of 2. At 16 hours post-infection, cells were fixed with 4% paraformaldehyde in PBS overnight at 4oC and permeabilized with 0.1 % Triton X-100 for 15 min at room temperature. Subsequently, the cells were subjected to immunofluorescence analysis for the detection of N and nsp1 proteins. For the detection of N protein, an affinity-purified rabbit anti-SARS-CoV N polyclonal antibody (Harcourt et al., 2020) was used as the primary antibody, followed by Alexa Fluor 488-conjugated secondary antibody (Invitrogen) and DAPI counterstaining, using SlowFade Gold antifade reagent (Invitrogen). The images for SARS-CoV-2 N protein were collected using an Olympus BX53 microscope with a 40X objective lens at UTMB and processed with ImageJ (NIH) software. For the detection of nsp1 protein, the anti-SARS-CoV-1 nsp1 antibody was used as the primary antibody, followed by Alexa Fluor 594-conjugated secondary antibody (Invitrogen) and DAPI counterstaining. The slides were mounted with VECTASHIELD Antifade Mounting Medium (Vector Laboratories). The images for SARS-CoV-2 nsp1 protein were collected using a Zeiss LSM 880 confocal microscope using a Plan-Apo 63X 1.40NA oil immersion lens at the UTMB Optical Microscopy Core. For extended figure 5a, cells were plated on ibidi slides, infected as described in ‘Cells and viruses’ at an MOI of 3 or left uninfected and at 8hpi washed once with PBS, fixed in 3% paraformaldehyde for 20 min, washed in PBS, permeabilized with 0.5% Triton X-100 in PBS for 2 min, and then blocked with 2% FBS in PBS for 30 min. Immunostaining was performed with rabbit anti-SARS-CoV-2 serum (Yahalom-Ronen et al., 2020) at a 1:200 dilution. Cells were washed and labeled with anti-rabbit FITC conjugated antibody at a 1:200 dilution and with DAPI (4′,6-diamidino-2-phenylindole) at a 1:200 dilution. Imaging was performed on a Zeiss AxioObserver Z1 wide-field microscope using a ×40 objective and Axiocam 506 mono camera.

### Northern blot analysis

Northern blot analysis was performed using total intracellular RNAs, as described previously (Kamitani et al., 2006). A digoxigenin (DIG)-labeled rLuc riboprobe was used to detect the Renilla luciferase gene, as described previously (Kamitani et al., 2006). A DIG-labeled random-primed probe, corresponding to nucleotides 28,999 to 29,573 of the SARS-CoV-2 genome (Kamitani et al., 2006; Xie et al., 2020), was used to detect SARS-CoV-2 mRNAs and visualized by DIG luminescent detection kit (Roche, Indianapolis, IN), according to the manufacturer’s protocol.

### Preparation of ribosome profiling and RNA-seq samples

SARS-CoV-2-infected cells were collected at 4 and 7 hpi. For RNA-seq, cells were washed with PBS and then collected with Tri-Reagent (Sigma-Aldrich). Total RNA was extracted and poly-A selection was performed using Dynabeads mRNA DIRECT Purification Kit (Invitrogen). The mRNA sample was subjected to DNase I treatment and 3′ dephosphorylation using FastAP Thermosensitive Alkaline Phosphatase (Thermo Scientific) and T4 polynucleotide kinase (New England Biolabs) followed by 3′ adaptor ligation using T4 ligase (New England Biolabs). The ligated products were used for reverse transcription with SSIII (Invitrogen) for first-strand cDNA synthesis. The cDNA products were 3′-ligated with a second adaptor using T4 ligase and amplified for 8 cycles in a PCR for final library products of 200–300 bp. For ribosome profiling libraries, cells were treated with 100 μg ml−1 cycloheximide for 1 min. Cells were then placed on ice, washed twice with PBS containing 100 μg ml−1 cycloheximide, scraped from 10-cm plates, pelleted and lysed with lysis buffer (1% triton in 20 mM Tris-HCl 7.5, 150 mM NaCl, 5 mM MgCl_2_, 1 mM dithiothreitol supplemented with 10 U/ml Turbo DNase and 100 μg/ml cycloheximide). After lysis, samples stood on ice for 2 h and subsequent ribosome profiling library generation was performed, as previously described(Finkel et al., 2020). In brief, cell lysate was treated with RNase I for 45 min at room temperature followed by SUPERase-In quenching. Sample was loaded on sucrose solution (34% sucrose, 20 mM Tris-HCl 7.5, 150 mM NaCl, 5 mM MgCl_2_, 1 mM dithiothreitol and 100 μg/ml cycloheximide) and spun for 1 h at 100,000 rpm using TLA-110 rotor (Beckman) at 4 °C. The pellet was collected using TRI reagent and the RNA was collected using chloroform phase separation. For size selection, 10 μg of total RNA was loaded into 15% TBE-UREA gel for 65 min, and 28–34-nt footprints were excised using 28-nt and 34-nt flanking RNA oligonucleotides, followed by RNA extraction and ribosome profiling library construction as previously described.

### Sequence alignment, metagene analysis, and differential gene expression analysis

Sequencing reads were aligned as previously described(Finkel et al., 2021b). In brief, linker (CTGTAGGCACCATCAAT for FP libraries and AGATCGGAAGAGCACACGT for mRNA libraries) and poly-A sequences were removed. Next, reads were aligned to a reference containing human rRNA and tRNA, and all reads that were successfully aligned were filtered out, and the remaining reads were aligned to the hg19 and to the SARS-CoV-2 genome (Genebank NC_045512.2). Alignment was performed using Bowtie v.1.1.229 with a maximum of two mismatches per read. Reads that were not aligned to the genome were aligned to the transcriptome (using the known canonical isoform UCSC gene annotations) and to SARS-CoV-2 junctions that were recently annotated(Kim et al., 2020). The aligned position on the genome was determined as the 5′ position of RNA-seq reads, and for ribosome profiling reads the p-site of the ribosome was calculated according to read length using the offset from the 5′ end of the reads that was calculated from canonical cellular ORFs. The offsets used were +12 for reads that were 28 or 29 bp and +13 for reads that were 30–33 bp. Footprint reads that were of other lengths were discarded. For the metagene analysis only genes with CDS length of at least 300 nucleotides, UTRs of at least 50 nucleotides and more than 50 reads in the analyzed window around the start or the stop codon were used. For each gene, reads were normalized to the position with the maximal number of reads in that gene, and then averaged. Differential expression analysis was done using DESeq2 (Love et al., 2014). ISG enrichment was calculated using a hypergeometric test, using a *p* value threshold of 0.001.

### Gene filtering, quantification and normalization

RNA-seq read coverage on CDS was normalized to RPKM to normalize for CDS length and sequencing depth. For analysis in which host and viral gene expression were compared (Fig. 4 and extended data figure 7), RPKM was normalized based on the number of reads uniquely aligned to the coding regions of both the host and the virus. For analysis that was focused on cellular gene expression, the RPKM was normalized based on the number of reads aligned to the rRNA and tRNA. This number was estimated by using the ratio of rRNA and tRNA reads to cellular aligned reads in a total RNA sequencing (without polyA selection). For footprint libraries, read coverage of cellular and viral genes was normalized to RPKM based on the number of reads aligned uniquely to the host total CDS aligned ribosome profiling host reads. Because the viral subgenomic transcripts are widely overlapping, RNA-seq RPKM levels of viral genes were computed with deconvolution as was previously described (Irigoyen et al., 2016; Kim et al., 2020). First, from the RPKM values of each subgenomic transcript, the RPKM of the upstream subgenomic (or the RPKM of ORF1b for the first subgenomic) transcript was subtracted. Then, for subgenomic transcripts, leader–body junctions were quantified on the basis of the number of uniquely mapped reads that span each canonical junction using STAR 2.5.3a aligner(Dobin et al., 2013). Finally, on the basis of the correlation between the deconvoluted RPKM and junction abundance of the subgenomic RNAs, the RPKM levels of all viral RNAs (including the genomic RNA) were estimated. Viral and host gene translation efficiency was calculated as the ratio between footprint RPKM and RNA-Seq RPKM. For cellular gene quantification, the number of reads aligned to the CDS of each gene in each replicate had to be greater than 10 reads.

### Quantification of intronic reads

To avoid biases from intron read count, first reads which are aligned to exons were excluded. Then the counting of reads for the introns was as described above for the exons, with intron annotations based on the known canonical isoform UCSC gene annotations. In addition, genes with a low number of reads (<10 on the exons and <2 on the introns) were omitted. The number of reads on exons and introns was normalized by the total length of the exons and introns, respectively, to get a quantification proportional to the number of molecules. Finally, the normalized number of reads on introns was calculated as a percentage of the normalized number of reads on exons. Genes with an intronic to exonic ratio of >1 were excluded from the analysis. Statistical significance (Fig. 5h) was tested using a paired t-test on the log values of the percentage.

### Ruxolitinib Assay

Calu3 cells were infected as described above with 0.01 MOI of SARS-CoV-2 wt or SARS-CoV-2 mut. After 1 h at 37°C, the cells were washed as detailed above and treated with 4μM Ruxolitinib (InvivoGen). Fresh Ruxolitinib was added at 24hpi.

### Protein synthesis measurement using OPP

The OPP assay (OPP, Thermo Fisher Scientific) was carried out following the manufacturer’s instructions. In brief, cells were collected following treatment with 10 μM OPP for 30 min at 37 °C. The cells were then fixed for 15 min in 3.7% formaldehyde, and permeabilized in 0.1% Triton X-100 for 15 min. OPP was then fluorescently labelled by a 30-min incubation in Click-iT Plus OPP reaction cocktail with Alexa Fluor594 picolyl azide (Thermo Fisher Scientific). Cells were analyzed using BD LSRII flow cytometer. The decrease in translation levels was calculated according to the median Alexa 594 fluorescence intensity between the 7 hpi and uninfected samples.

### Subcellular Fractionation

293T cells transfected with a plasmid expressing Nsp1-WT, nsp1-ΔRB, nsp1-CD, or GFP were washed in PBS at 24 h post-transfection, trypsinized and resuspended in cold PBS. A fraction containing 10% of the cells was then transferred to a new tube and RNA was extracted in Tri-reagent to obtain whole cellular extract. Remaining cells were pelleted for 5 min at 300g. Cells were resuspended in 150 μl fractionation buffer A (15 mM Tris-HCl pH 8.0, 15 mM NaCl, 60 mM KCl, 1 mM EDTA pH 8.0, 0.5 mM EGTA pH 8.0, 0.5 mM spermidine, and 10 U ml−1 RNase inhibitor), and 150 μl 2x lysis buffer (15 mM Tris-HCl pH 8.0, 15 mM NaCl, 60 mM KCl, 1 mM EDTA pH 8.0, 0.5 mM EGTA pH 8.0, 0.5 mM spermidine,10 U ml−1 RNase inhibitor and 0.5% NP-40) was added followed by 10 min incubation on ice. The extract was pelleted for 5 min at 400g and the supernatant containing the cytoplasmic fraction was removed to a new tube. This was centrifuged again at 500g for 1 min, the supernatant was transferred to a new tube and RNA was extracted with Tri-reagent. Thenuclear pellet was resuspended in 1 ml RLN buffer (50 mM Tris-HCl pH 8.0, 140 mM NaCl, 1.5 mM MgCl_2_, 0.5% NP-40, 10 mM EDTA and 10 U/ml RNase inhibitor) and incubated on ice for 5 min. The nuclear fraction was then pelleted for 5 min at 500g, the supernatant was removed and RNA was extracted from the pellet with Tri-reagent. RNA-seq libraries were then prepared from all three fractions as described above.

### Subcellular Fractionation analysis

RNA-seq reads from total, nuclear and cytosolic fractions were aligned to the human and viral reference as described above. Human gene read counts were adjusted to RPKM as described above, and then converted to transcripts per million (TPM) by normalizing to the sum of RPKM in each sample, so that the expression levels in each sample sum up to the same value.

A list of 2203 average-expressed genes was defined. These genes were genes with 15 or more reads across all samples and with a sum of TPM values in the total RNA samples across replicates within quantiles 0.4 and 0.9. On the basis of this list, for each replicate, a linear regression model was calculated of the total fraction as a linear combination of the cytosolic and the nuclear fractions. The regression coefficients were used to normalize the cytosolic and nuclear TPM values to obtain absolute localization values(Carlevaro-Fita et al., 2019). To correct for changes in total mRNA levels, the absolute values were further scaled by a factor calculated from total RNA-seq as described in ‘Gene filtering, quantification and RPKM normalization. To statistically compare the effect of infection on nuclear–cytosolic distribution of mRNAs from different clusters, P values were calculated from the interaction term in a linear model.

### RNA labelling for SLAM-seq

For metabolic RNA labelling of 293T transfected with a plasmid or Calu3 infected cells, growth medium was replaced with medium containing 4sU (T4509, Sigma) 24 h post-transfection and 3 hpi at a final concentration of 200 μM (a concentration that did not induce substantial cell cytotoxicity at 3 h labelling). Cells were collected with Tri-reagent at 2, 3, or 4 h after labelling for transfected cells or at 1, 2, or 3 h after labelling for infected cells (corresponding to 4, 5, and 6 hpi). RNA was extracted under reducing conditions and treated with iodoacetamide (A3221, Sigma) as previously described(Herzog et al., 2017). RNA-seq libraries were prepared and sequenced as described above.

### SLAM-seq data analysis and half-life calculation

Alignment of SLAM-seq reads was performed using STAR(Dobin et al., 2013; Herzog et al., 2017), with parameters that were previously described(Finkel et al., 2021b). First, reads were aligned to a reference containing human rRNA and tRNA, and all reads that were successfully aligned were filtered out. The remaining reads were aligned to a reference of the human and the virus as described above. Reads mapped to the virus were discarded and reads mapped to the human were used in the next steps. Output.bam files from STAR were used as input for the GRAND-SLAM analysis(Jürges et al., 2018) with default parameters and with trimming of 5 nucleotides in the 5′ and 3′ends of each read. Infected and uninfected samples were analyzed in separate runs. Each one of the runs also included an unlabeled sample (no 4sU) that was used for estimating the linear model of the background mutations. The output of GRAND-SLAM, i.e., the estimated ratio of newly synthesized out of total molecules for each gene (new to total ratio (NTR)), was used to calculate the half-life as described below. The old transcript fraction for each gene in each sample is 1 – NTR; this number reflects the pre-existing mRNA molecules (not labelled) and these values were used for half-life estimation of cellular genes. The half-life of each gene was calculated by linear regression of the log values of the calculated old transcript fraction. Estimated variance of the values as calculated by GRAND-SLAM were used as weights in the linear regression. The regression coefficient λ was converted to half-life as −log(2)/λ. For further analysis, only genes for which the P value in the regression was < 0.1 and the adjusted r2 > 0.7 were used.

### Plasmids and cloning

pLVX-EF1alpha-SARS-CoV-2-nsp1-2XStrep-IRES-Puro (nsp1-WT) was kindly provided by N. Krogan. The mutations to generate nsp1-ΔRB and nsp1-CD were introduced by amplifying the nsp1-WT plasmid so that it became a linear fragment containing the desired mutations (primers 1-4 listed in Supplementary Table 1).

For the reporter assay, reporter plasmids containing both the CoV-2 sub-genomic leader and a human control 5’UTR were used as described in(Finkel et al., 2021a). In order to express Nsp1-WT, nsp1-ΔRB, and nsp1-CD under the SARS-CoV-2 leader, the nsp1 cassette, including the twin strep II tag at the C-terminus of the protein, was lifted from their respective plasmids and inserted immediately downstream of the SARS-CoV-2 subgenomic leader sequence on the plasmid used in the reporter assay in place of the GFP cassette, using restriction-free cloning (primers 5 & 6 Supplementary Table 1).

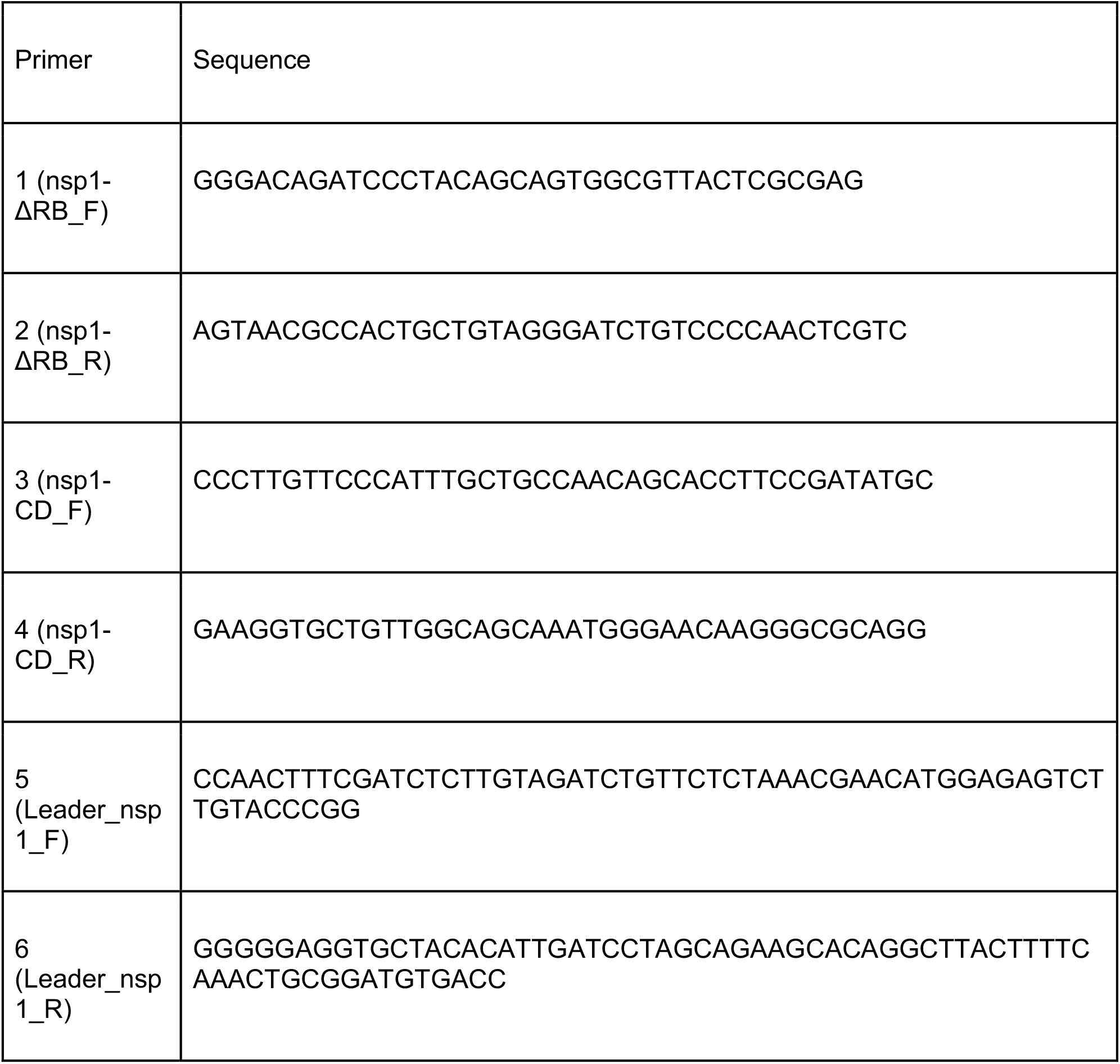

### Reporter assay

293T cells were transfected using JetPEI (Polyplus-transfection) following the manufacturer’s instructions. Twenty-four h after transfection, cells were assayed for reporter expression by flow cytometry on a BD Accuri C6 flow cytometer. In parallel, cells were assayed for expression of NSP1 and reporter mRNA levels as detailed below.

### Flow cytometry analysis of strep tag

The expression of NSP1 was verified by staining of the fused strep-tag, followed by flow cytometry. Cells were fixed in 4% paraformaldehyde, permeabilized in 0.1% Triton X100, and stained using Strep-TactinXT DY-649 (IBA-lifesciences). Flow cytometry analysis was performed on BD Accuri C6 and analysed on FlowJo. Normalization to the mode is presented in the histograms.

### Quantitative real-time PCR analysis

Total RNA was extracted using Direct-zol RNA Miniprep Kit (Zymo Research) following the manufacturer’s instructions. cDNA was prepared using qScript FLEX cDNA Synthesis Kit with random primers (Quanta Biosciences) following the manufacturer’s instructions.

Real-time PCR was performed using the SYBR Green PCR master-mix (ABI) on the QuantStudio 12K Flex (ABI) with the following primers (forward, reverse): GFP (TGACCCTGAAGTTCATCTGC, GAAGTCGTGCTGCTTCATGT); and 18S (CTCAACACGGGAAACCTCAC, CGCTCCACCAACTAAGAACG).

GFP mRNA levels were calculated relative to 18S rRNA.

### Ribosome pelleting and Western blot analysis

293T cells were transfected with Nsp1-WT, nsp1-ΔRB, or nsp1-CD. At 24 h post-transfection cells were lysed in lysis buffer (1% triton in 20 mM Tris-HCl, pH 7.5, 150 mM NaCl, 5 mM MgCl_2_, 1 mM dithiothreitol and 10 U ml−1 Turbo DNase). The lysate was loaded on sucrose solution (34% sucrose, 20 mM Tris-HCl 7.5, 150 mM NaCl, 5 mM MgCl_2_, and 1 mM dithiothreitol) and spun for 1 h at 100,000 rpm using TLA-110 rotor (Beckman) at 4 °C. The ribosome pellet was suspended in RIPA buffer and the amount of protein was quantified from both the ribosomal fraction as well as the whole lysate by Pierce^™^BCA Protein Assay (Thermo Fisher Scientific). Samples were then separated by 4-12% polyacrylamide Bis-tris gel electrophoresis (Invitrogen), blotted onto a nitrocellulose membrane and immunoblotted with Streptactin-HRP (Bio-rad), ɑRPS6 (Cell signaling Technology, 2317), and ɑGAPDH (Cell signaling Technology, 2118). Secondary antibodies used were Goat anti-rabbit or Goat anti-mouse (IRDye 800CW or IRDye 680RD, Li-Cor). Reactive bands were detected by Odyssey CLx infrared imaging system (Li-Cor) and quantified using Fiji (ImageJ) as detailed in https://imagej.nih.gov/nih-image/manual/tech.html. For figure 3, western blot analysis was performed, as described previously (Kamitani et al., 2006). An affinity-purified rabbit anti-SARS-CoV-1 nsp1 polyclonal antibody (Narayanan et al., 2015) was used as the primary antibody, and an HRP-linked anti-rabbit IgG (Cell Signaling Technology) was used as the secondary antibody.

### Graphics

Figure 1a and extended figure 2c were created using BioRender. Fluorescence-activated cell sorting figures were created with FlowJo and the rest of the figures were drawn with ggplot2 in R.

